# Neural circuitry for maternal oxytocin release induced by infant cries

**DOI:** 10.1101/2021.03.25.436883

**Authors:** Silvana Valtcheva, Habon A. Issa, Chloe J. Bair-Marshall, Kathleen A. Martin, Kanghoon Jung, Yiyao Zhang, Hyung-Bae Kwon, Robert C. Froemke

## Abstract

Oxytocin is a neuropeptide important for maternal physiology and childcare, including parturition and milk ejection during nursing^1–8^. Suckling triggers oxytocin release, but other sensory cues- specifically infant cries- can elevate oxytocin levels in new human mothers^9–11^ indicating that cries can activate hypothalamic oxytocin neurons. Here we describe a neural circuit routing auditory information about infant vocalizations to mouse oxytocin neurons. We performed in vivo electrophysiological recordings and photometry from identified oxytocin neurons in awake maternal mice presented with pup calls. We found that oxytocin neurons responded to pup vocalizations, but not pure tones, via input from the posterior intralaminar thalamus, and repetitive thalamic stimulation induced lasting disinhibition of oxytocin neurons. This circuit gates central oxytocin release and maternal behavior in response to calls, providing a mechanism for the integration of sensory cues from the offspring in maternal endocrine networks to ensure modulation of brain state for efficient parenting.

## Introduction

Parenting behaviors emerge from complex neural circuits conferring sensitivity to infant needs to ensure survival of the species. One important molecular signal for the maternal brain is oxytocin, a nine amino acid peptide produced mainly in the paraventricular nucleus (PVN) and supraoptic nucleus of the hypothalamus^1–4^. Peripheral oxytocin is important for milk ejection during nursing and uterine contractions during labor, whereas oxytocin release in the central nervous system is involved in a wide range of behaviors including reproduction, parental care and pair bonding^5–8^.

In humans, baby cries are the most powerful signal of infant distress, and most nursing mothers respond to cries with increased hypothalamic activity, elevations of plasma oxytocin, comforting behaviors towards the infant, and occasional milk ejection^9–11^. Postpartum conditions might relate to lower oxytocin levels and decreased sensitivity to infant cries^10, 12^, underscoring the importance of understanding the mechanisms by which acoustic stimuli from the offspring can activate oxytocin neurons. PVN oxytocin cell receive projections from many brain areas, but it is unknown which specific pathways relay auditory information and what synaptic mechanisms might mediate auditory responses and oxytocin release from oxytocin cells^3, 13–16^.

## Results

We made cell-attached and whole-cell recordings from PVN oxytocin neurons (ChR2+; OT+) and other optically-unresponsive PVN neurons (ChR2-; OT-) in awake head-fixed mouse dams by using channelrhodopsin2-assisted patching (**Fig. 1a-c**). Oxytocin neurons were identified by reliable spiking responses to brief pulses of blue light with comparable latencies to previous reports^16, 17^, and had baseline firing rates similar to those formerly described^16–20^ (**Extended Data Fig. 1 and 2a**).

**Figure 1.**
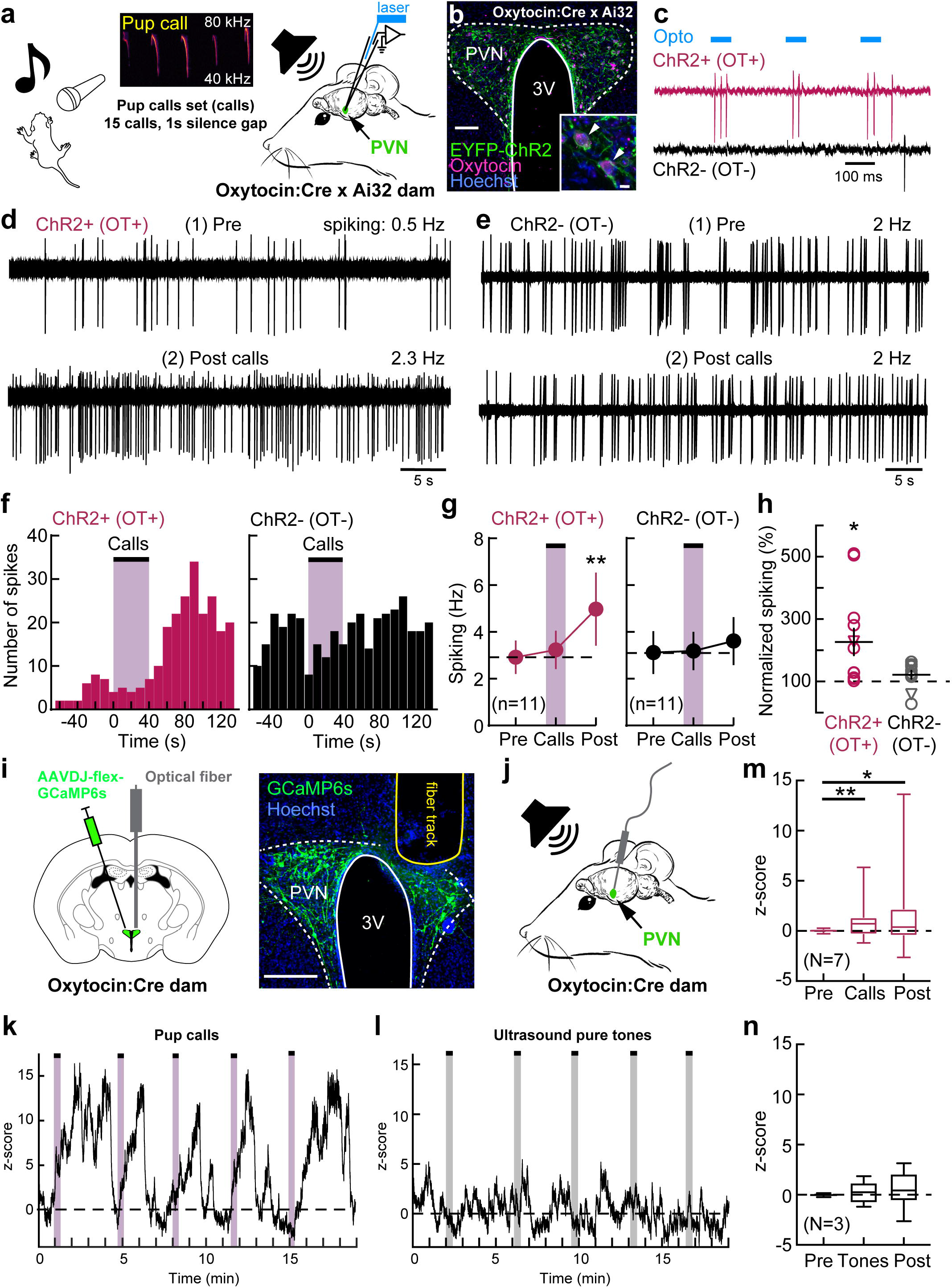
Delayed persistent activation of dam oxytocin neurons by pup vocalizations. (a) In vivo cell-attached/whole-cell channelrhodopsin2-assisted recordings and playback of pup calls. (b) Oxytocin cells expressed EYFP-ChR2. Scale, 100 µm (inset, 10 µm). 3V, third ventricle. (c) In vivo optogenetic identification of ChR2+ (OT+) neurons with reliable spiking to blue light pulses (‘Opto’). (**d, e**) Cell-attached recordings of ChR2+ (OT+), ChR2- (OT-) neurons before pup call onset (1, ‘Pre’), and 80 sec after call onset (2, ‘Post’). (f) Peristimulus time histograms for ChR2+ (OT+) and ChR2- (OT-) neurons. Bins: 10 s. (g) Firing rate of ChR2+ (OT+; n=11 neurons, N=6 dams; p=0.0014, Friedman test) and ChR2- (OT-; n=11, N=5; p=0.07) neurons. (h) Change in firing rate of ChR2+ (OT+; p=0.01, one-sample two-tailed Student’s t-test) but not ChR2- (OT-; p=0.12) neurons. Cell-attached (circles) and whole-cell (triangles) recordings. (i) Injection of AAVDJ-CAG-FLEx-GCaMP6s in PVN of Oxytocin:Cre dams and fiber implantation. Scale, 200 µm. (j) Fiber photometry in awake head-fixed dams and pup calls playback via ultrasonic speaker. (**k-n**) Oxytocin neurons responded to pup calls but not pure tones. Example timelines of responses to five sets of pup calls (**k**) or ultrasound pure tones (**l**). Z-scores of fluorescence activity preceding onset of pup calls or tones (‘Pre’), during auditory stimulus (‘Calls/Tones’) and after (‘Post’) (**m**; N=7, Pre vs Calls: p=0.0048; Pre vs Post: p=0.0179; **n**; N=3, Pre vs Calls: p=0.2119; Pre vs Post: p=0.3394; Wilcoxon matched-pairs signed rank one-tailed test and correction for multiple comparisons). Data reported as mean±SEM or median±min to max (**m and n**). *p<0.05, **p<0.01.

**Figure 2.**
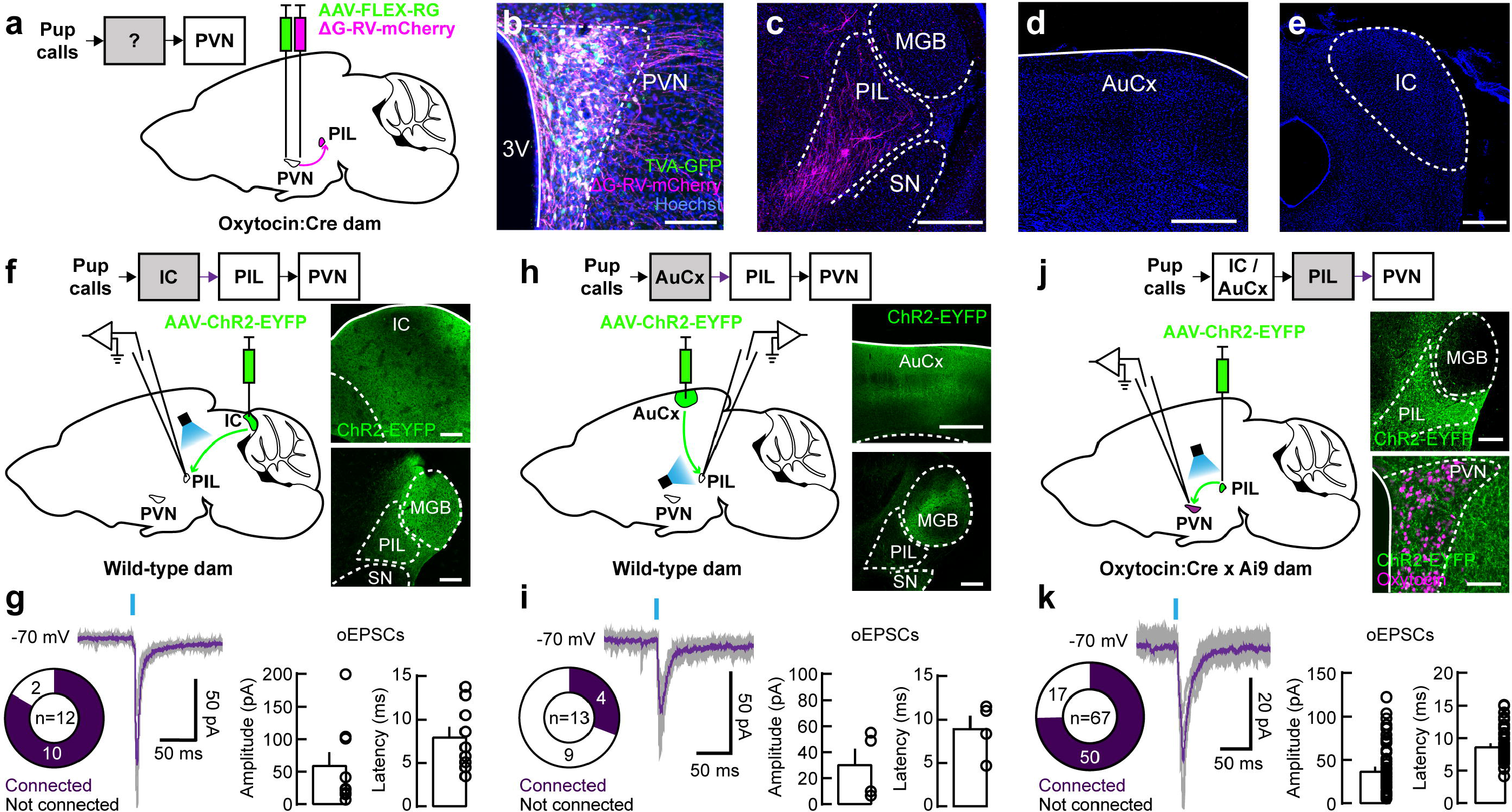
PVN oxytocin neurons receive projections from the auditory thalamus. (**a-e**) Rabies virus tracing. (**a**) Injection of helper virus AAV2-EF1a-FLEX-TVA-GFP followed by *Cre*-inducible, retrograde pseudotyped monosynaptic rabies virus SADΔG- mCherry. (**b**) Starter oxytocin neurons in PVN expressing TVA-GFP and ΔG-RV-mCherry. Scale, 100 µm. (**c-e**) Retrograde rabies infection and mCherry staining was absent in MGB (**c**), AuCx (**d**), and IC (**e**). Dense rabies-infected field and robust expression of mCherry was found in PIL (**c**; N=4 dams). Scale, 500 µm. AuCx, auditory cortex; IC, inferior colliculus; MGB, medial geniculate body of the thalamus; PIL, posterior intralaminar nucleus of the thalamus; SN, substantia nigra. (**f, g**) PIL receives reliable input from IC. (**f**; left) Injection of AAV1-hSyn-hChR2(H134R)- EYFP in IC of wild-type dams and whole-cell voltage-clamp recordings in PIL slices. (Right) ChR2-EYFP expression in IC and IC projections in MGB and PIL. Scale, 200 µm. (**g**) Characterization of oEPSCs in PIL neurons triggered by optogenetic stimulation of IC axons (10/12 connected neurons, N=5). (**h, i**) PIL receives sparser input from AuCx. (**h**; left) Injection of AAV1-hSyn- hChR2(H134R)-EYFP in AuCx of wild-type dams and whole-cell voltage-clamp recordings in PIL slices. (Right) ChR2-EYFP expression in AuCx and dense AuCx projections in MGB but sparser in PIL. Scales, 500 µm and 200 µm, respectively. (**i**) oEPSCs in PIL neurons triggered by optogenetic stimulation of AuCx axons (4/13 connected neurons, N=3). (**j, k**) PVN oxytocin neurons receive reliable input from PIL. (**j**) Injection of AAV1-hSyn- hChR2(H134R)-EYFP in PIL of Oxytocin:Cre x Ai9 dams and whole-cell voltage-clamp recordings from oxytocin neurons (tdTomato+). (**j**; left) ChR2-EYFP expression in PIL and PIL projections in PVN. Scales, 200 µm and 100 µm, respectively. (**k**) oEPSCs in oxytocin neurons triggered by optogenetic stimulation of PIL axons (50/67 connected, N=21). Data reported as mean±SEM.

To investigate whether ChR2+ (OT+) and/or ChR2- (OT-) neurons were activated by pup distress vocalizations, we measured 1-2 min of baseline spiking (‘Pre’) followed by repetitive presentation of a set of 15-18 distress calls recorded from isolated pups (‘Calls’, each one- second call followed by one second of silence for a total duration of ∼30-40 s), and assessed changes in ongoing activity thereafter for the duration of the recordings (‘Post’). We found that pup call presentation increased the firing rates of ChR2+ (OT+) neurons (Pre: 2.9±0.7 Hz, Post: 5.0±1.6 Hz, p=0.001, n=11 cells from N=6 dams), but not ChR2- (OT-) neurons (Pre: 3.1±0.9 Hz, Post: 3.6±1.0 Hz, p=0.07, n=11, N=5; **Fig. 1d-h, Extended Data Fig. 2b-f**). This increase in spiking of ChR2+ (OT+) neurons was not observed during call presentation itself (**Extended Data Fig. 2g-j**). To examine if the population-level activity was also enhanced after pup calls, we performed fiber photometry of oxytocin neuronal responses in awake head-fixed Oxytocin:Cre dams injected with AAVDJ-CAG-FLEx-GCaMP6s in PVN and implanted with an optical fiber above PVN (**Fig. 1i**). We measured changes in GCaMP6s signals in oxytocin neurons while playing pup distress calls and observed substantial responses following calls (Pre vs Calls: p=0.0048; Pre vs Post: p=0.0179, N=7 dams; **Fig. 1j, k and m, Extended Data Fig. 3a, i and j**) and this increase in activity was significant as early as at 25 sec from call onset (p=0.0134). The sustained increase in oxytocin neuron firing was specific to pup calls, as PVN neurons did not respond to pure tones, adult calls or frequency-modulated down-sweeps which retained key spectrotemporal characteristics of pup vocalizations (**Fig. 1l and n, Extended Data Fig. 3c, e, g, h and j**). We also did not observe significant activation of oxytocin cells in virgins consistent with our previous findings^17^ (**Extended Data Fig. 3b, d, f and j**). These results show that pup vocalizations can activate PVN oxytocin neurons, but not with stimulus-locked responses such as those typically observed in the central auditory system including the auditory cortex^21–24^. Instead, oxytocin cells require more prolonged stimulus periods to increase activity, perhaps as might naturally occur before and during episodes of maternal care.

**Figure 3.**
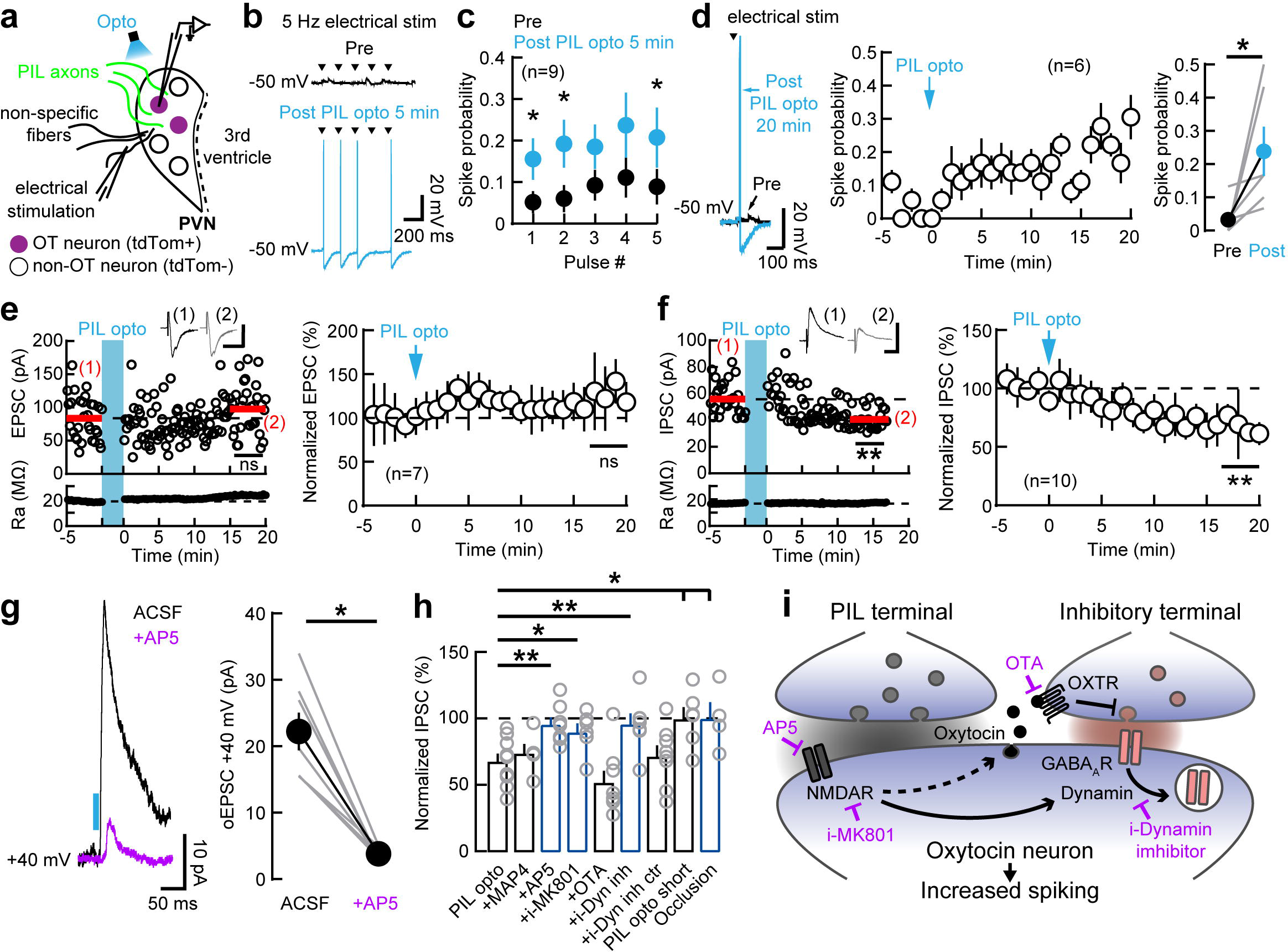
Activation of PIL inputs to PVN decreases inhibition in oxytocin neurons via postsynaptic NMDARs and dynamin-dependent internalization of GABA_A_Rs. (a) Whole-cell recordings from tdTomato+ oxytocin neurons in PVN slices, optogenetic stimulation of PIL axons and placement of the extracellular stimulation electrode. (**b and c**) Short-term increase in spiking probability of oxytocin neurons following PIL opto. Whole-cell current-clamp recording showing improved spiking output of oxytocin neuron in response to 5 Hz electrical stimulation at 5 min after PIL opto (‘Post’) compared to baseline activity (‘Pre’): example trace (**b**) and summary (**c**; n=9 neurons, p<0.05 Wilcoxon matched- pairs signed rank two-tailed test). (d) Long-term increase in spiking probability of oxytocin neurons following PIL opto. Left, example trace showing improved spiking output of oxytocin neuron at 20 min after PIL opto in response to a single pulse of electrical stimulation. Middle, timeline of long-term spiking probability after PIL opto (n=6). Right, spiking probability was significantly increased at 20 min after PIL opto (n=6, p=0.03, Wilcoxon). (**e, f**) Repeated optogenetic stimulation of PIL axons (‘PIL opto’) induced iLTD in oxytocin neurons. (**e**) No change in EPSC magnitude after PIL opto; example cell (left; p=0.15, Mann- Whitney two-tailed test; scale, 20 ms and 50 pA) and summary (right; n=7, p=0.14, one- sample two-tailed Student’s t-test). (**f**) Decreased IPSC magnitudes after PIL opto; example cell (left; p<0.0001, Mann-Whitney; scale, 20 ms and 100 pA) and summary (right; n=10, p=0.0003, one-sample Student’s t-test). (g) NMDAR-dependent currents in oxytocin cells in response to a single pulse of optogenetic stimulation of PIL axons (n=7, p=0.02, Wilcoxon). (h) iLTD is blocked by AP5 (p=0.0021, Mann-Whitney), i-MK801 (p=0.02), i-Dynamin inhibitor (p=0.0068) and occluded by presentation of pup calls (“Occlusion”, p=0.0140), but not by MAP4 (p=0.37), OTA (p=0.08); PIL opto vs PIL opto short (p=0.02), or PIL opto vs - Dynamin inhibitor control (p=0.8968). (i) Possible mechanisms for postsynaptic NMDAR-dependent decrease in inhibition. OXTR, oxytocin receptor. Data reported as mean±SEM (**c, d right, g, h**) or as mean±SD (**d middle, e, f**). *p<0.05, **p<0.01, ns: not significant.

What projections relay information about pup calls to PVN oxytocin neurons? We used *Cre*- inducible, retrograde pseudotyped monosynaptic rabies virus to determine the inputs to oxytocin cells (**Fig. 2a and b**). Consistent with the absence of phase-locked responses to auditory stimuli in PVN neurons, most auditory areas did not send direct projections to the PVN. We did not detect retrograde rabies infection and mCherry staining in the medial geniculate body of the thalamus (MGB; **Fig. 2c**), auditory cortex (AuCx; **Fig. 2d**), or the inferior colliculus (IC; **Fig. 2e**). However, we did observe dense and reliable rabies-based mCherry staining in the posterior intralaminar nucleus of the thalamus (PIL; **Fig. 2c**). PIL contributed 100% to the auditory inputs in PVN oxytocin cells versus 0% for inputs from MGB, AuCx and IC. PIL is part of the non-lemniscal auditory pathway which projects to the PVN^16, 25^, and it is also implicated in maternal and social behaviors^25–27^.

We found that PIL is activated by pup calls. Even though firing rates of PIL during pup calls and ultrasound pure tones are similar (p=0.33), pure tones evoked only a transient activity for less than 50 ms duration, as previously described^27^, compared to 1 second or more during pup calls (**Extended Data Fig. 4**). We then explored if PIL receives functional synaptic inputs from the IC and AuCx, two major auditory areas that respond to pup calls. We tested the strength of these synaptic connections using channelrhodopsin2-assisted circuit mapping. We injected AAV1-hSyn-hChR2(H134R)-EYFP in the IC (**Fig. 2f and g**) or AuCx (**Fig. 2h and i**) of wild-type dams, and performed whole-cell voltage-clamp recordings from PIL neurons in acute brain slices while optogenetically stimulating axons from either IC or AuCx. We found that PIL receives synaptic inputs from both IC and AuCx with a bigger proportion of PIL neurons receiving inputs from IC (p=0.0154, two-tailed Fisher’s test; **Fig. 2g, i**). These findings are similar to those in other neurons in the paralaminar thalamus, which are more efficiently driven by inputs from IC than AuCx^28^.

**Figure 4.**
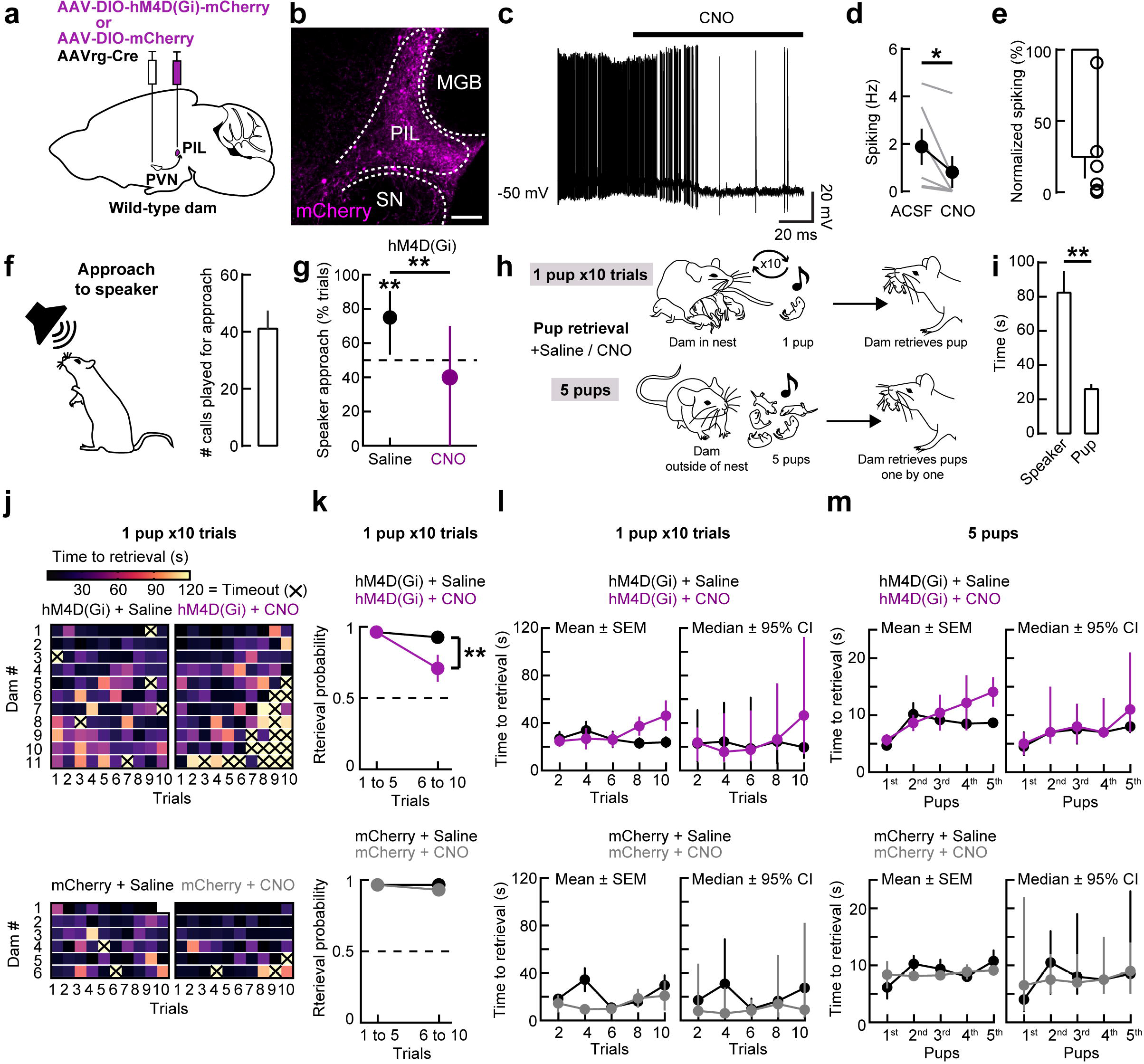
Chemogenetic inhibition of PIL projections to PVN impairs detection of pup calls and pup retrieval behavior. (a) Injection of AAVrg-ENN.AAV.hSyn.Cre.WPRE.hGH in PVN and AAV8-hSyn-DIO- hM4D(Gi)-mCherry or AAV8-hSyn-DIO-mCherry in PIL. (b) mCherry expression in PVN-projecting PIL neurons. Scale, 200 µm. (**c-e**) Bath-application of CNO (1 µM) reduced firing rate in PIL slices from dams expressing hM4Di. Example whole-cell current-clamp recording from PIL neuron (**c**) and summary (**d**; n=6 neurons, p=0.03, Wilcoxon matched-pairs signed rank two-tailed test; **e**; p=0.003, one- sample two-tailed Student’s t-test). (**f-g**) Dams expressing hM4Di and injected with CNO had no preference for pup calls. (**f**) Schematic of speaker approach protocol and number of calls played for approach. (**g**) Approach probability (43 trials; N=4 dams; p=0.002, two-tailed Fisher’s test; two-tailed Binomial test, Saline: p=0.0054, CNO: p=0.1173). (**h-m**) Dams expressing hM4Di and injected with CNO had impaired pup retrieval behavior. (**h**) Pup retrieval protocols. (**i**) Latency to approach a speaker playing pup calls vs crying pup (p<0.0001, Mann-Whitney two-tailed test). (**j-l**) 1 pup 10 trials test: (**j**) Performance of all dams across trials. (**k**) Dams expressing hM4Di and injected with CNO retrieved with lower probability compared to saline controls at trials 6 to 10 (N=11, p=0.0055, Fisher’s test); there was no difference in dams expressing mCherry (N=6, p>0.9999, Fisher’s test). (**l**) No difference in time to retrieval for trials 6 to 10 (hM4Di: N=11, p>0.1042, Mann-Whitney; N=11; mCherry: N=6, p>0.3939, Mann-Whitney). (**m**) 5 pups test: No difference in time to retrieval (hM4Di: N=6 dams, p>0.0556, Mann-Whitney; mCherry: N=4, p>0.3142, Mann- Whitney). Data reported as mean±SEM or as median±95% CI (g, l and m). *p<0.05, **p<0.01.

We then examined the strengths of synaptic connections between PIL and PVN oxytocin neurons^16, 25^. We injected AAV1-hSyn-hChR2(H134R)-EYFP in PIL of Oxytocin:Cre x Ai9 dams, to perform in vitro whole-cell voltage-clamp recordings from identified oxytocin neurons (expressing tdTomato) in acute brain slices of PVN while optogenetically stimulating PIL terminals (**Fig. 2j**). We observed reliable oEPSCs in the majority of oxytocin cells indicating that oxytocin neurons receive connections from PIL with high probability (**Fig. 2k**). These monosynaptic connections were both excitatory^29^ and inhibitory and preferentially targeted parvocellular oxytocin neurons (**Extended Data Fig. 5a-h**). Consistent with our rabies virus tracing experiments, we did not observe axon terminals in PVN originating from IC or AuCx (**Extended Data Fig. 5i and j**). In summary, our findings describe a noncanonical auditory circuit that could relay pup vocalization signals from IC and AuCx via PIL to PVN oxytocin neurons.

**Figure 5.**
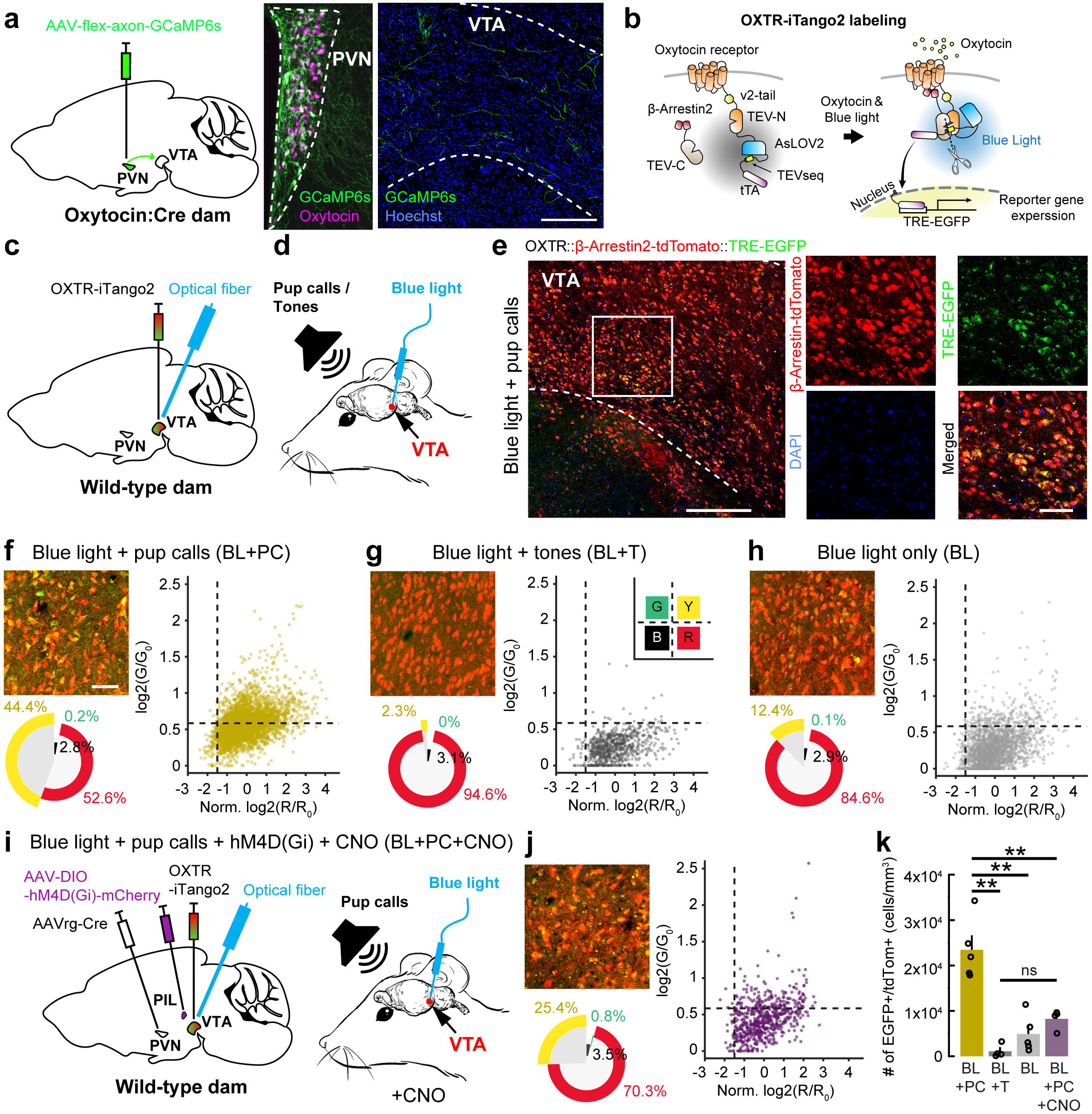
Pup calls trigger oxytocin release in the VTA via PIL-to-PVN pathway. (a) Injection of AAV5-hSynapsin1-FLEx-axon-GCaMP6s in the PVN of Oxytocin:Cre dams and GCaMP6s+ axons in the VTA. Scale, 200 µm. N=2 dams. (b) Schematic of *OXTR-iTango2* genetic strategy. (c) Injection of *OXTR-iTango2* viral constructs and fiber implantation in VTA. (d) Dams were exposed to either pup calls paired with blue light (‘Blue light+pup calls’; ‘BL+PC’), pure tones paired with blue light (‘Blue light+tones’; ‘BL+T’), or blue light alone (‘Blue light only’; ‘BL’). (e) Images of VTA neurons expressing *OXTR-iTango2* constructs for Blue light+pup calls (left; scale, 200 µm) and magnified images from the area marked by a square (right; scale, 50 µm). (**f-h**) Distribution patterns and percentage of *OXTR-iTango2* labeled neurons VTA of dams. Scatter plots of individual cells (black dashed lines indicate threshold): cells with no viral infection (tdTomato-/EGFP-; black, ‘B’); cells with only TRE-reporter signal (tdTomato-/EGFP+; green, ‘G’), cells with only β-Arrestin-tdTomato signal (tdTomato+/EGFP-; red, ‘R’); cells with both red and green signal (tdTomato+/EGFP+; yellow, ‘Y’). BL+PC (N=5; **f**), BL+T (N=4; **g**), and BL (N=5; **h**). Scale, 50 µm. (i) Injection of AAVrg-ENN.AAV.hSyn.Cre.WPRE.hGH in PVN, AAV8-hSyn-DIO- hM4D(Gi)-mCherry in PIL and *OXTR-iTango2* viral constructs in VTA together with fiber implantation in VTA. Dams were injected with CNO and exposed to pup calls paired with blue light (‘Blue light+pup calls+hM4D(Gi)+CNO’; ‘BL+PC+CNO’; N=4). (j) Distribution patterns and percentage of *OXTR-iTango2* labeled neurons VTA of dams expressing hM4D(Gi) in PVN-projecting PIL cells and injected with CNO. (k) Density quantification of *OXTR-iTango2* labeled cells. There was a significant amount of yellow (tdTomato+/EGFP+) cells in BL+PC compared to BL+T (p<0.0001, one-way ANOVA; p<0.0001, *post hoc* Bonferroni correction), BL (p=0.0001), or BL+PC+CNO (p=0.0008). There was no difference in the amount of yellow cells between BL+T, BL and BL+PC+CNO. Data reported as mean±SEM. **p<0.01, ns: not significant.

Persistent activation of oxytocin neurons following pup calls can be due to several mechanisms: 1) long-term increase in PIL activity following pup calls, 2) changes in intrinsic excitability, and 3) synaptic plasticity. We performed in vivo cell-attached recordings from PIL neurons in wild-type dams, and while we observed stimulus-evoked auditory responses in the PIL (**Extended Data Fig. 4**), pup calls did not trigger a long-term increase in spontaneous spiking (**Extended Data Fig. 6**). This makes unlikely that persistent activation of PVN oxytocin neurons resulted from increased PIL output following pup calls. We then asked if increased firing of PVN oxytocin neurons could be caused by changes of intrinsic excitability, or synaptic plasticity mechanisms. We performed whole-cell current-clamp recordings from oxytocin neurons in PVN slices and found that although PIL inputs to these cells occur with high probability (**Fig. 2k**), the synaptic connections are too weak to trigger postsynaptic spiking responses (**Extended Data Fig. 7a-c**). We then used a stimulation pattern emulating PIL responses in vivo during pup call playback by optogenetically stimulating PIL terminals in PVN (‘PIL opto’) at 30 Hz for one second and repeated this high-frequency activation every two seconds for three minutes. However, this stimulation pattern also failed to elicit suprathreshold spiking in oxytocin cells (**Extended Data Fig. 7d**), possibly explaining why oxytocin neurons are not activated during individual pup calls in vivo (**Extended Data Fig. 2h-k**). There were no changes of intrinsic properties of oxytocin neurons following PIL opto (**Extended Data Fig. 7e-i**), indicating that increased output of these cells in vivo is unlikely due to such mechanism.

We observed a rapid and enduring increase in evoked spike generation in oxytocin neurons following PIL opto (**Fig. 3a-d**), similar to the persistent activity of oxytocin cells we observed in vivo following pup calls. We asked if this repetitive PIL stimulation led to lasting changes in excitatory or inhibitory inputs onto oxytocin neurons. Reduction of inhibition could increase output of these cells as PVN is heavily innervated by inhibitory networks^30^. We monitored excitatory and inhibitory postsynaptic currents (EPSCs and IPSCs) evoked by an extracellular stimulation electrode, before and after PIL opto (**Fig. 3e and f**). While EPSCs were unaffected, the amplitudes of IPSCs were significantly decreased following PIL opto, without affecting PIL transmission itself (**Extended Data Fig. 8a and b**). Correspondingly, this long-term depression of inhibition (iLTD) decreased the IPSC/EPSC ratio for oxytocin neurons (**Extended Data Fig. 8c**). In contrast, emulating PIL responses in vivo during pure tones playback by brief optogenetic stimulation of PIL terminals in PVN (‘PIL opto short’) at 30 Hz for 100 ms repeated every two seconds for three minutes did not trigger iLTD (**Extended Data Fig. 8d-f**). These findings are in accordance with our in vivo data showing that sustained increase in firing of oxytocin neurons was specific to pup calls. In addition, iLTD was occluded in slices of dams which were exposed to pup calls playback immediately before brain slices preparation (‘Occlusion’; **Fig. 3h; Extended Data Fig. 8g-i**), suggesting that disinhibition of oxytocin cells via iLTD occurs following pup calls in vivo.

Long-term reduction in inhibitory transmission can be mediated by different mechanisms. We found that type-III metabotropic glutamate receptors (mGluRs), which can be located on GABAergic terminals^31^, were unlikely to be responsible for the enhanced oxytocin neuron firing after pup calls as the type-III mGluR antagonist MAP4 did not prevent iLTD induction (**Extended Data Fig. 9a**). Next, we explored if iLTD relies on pre- or postsynaptic NMDARs. Optogenetic stimulation of PIL fibers activated NMDAR receptors on oxytocin neurons as it elicited outward oEPSCs, which were blocked by the NMDAR antagonist AP5 (p=0.02; **Fig. 3g**). Furthermore, both bath-application of AP5, or intracellular delivery of use-dependent NMDAR blocker MK801 (i-MK801) via recording pipette abolished iLTD after PIL opto (**Extended Data Fig. 9b and c**). These results show that iLTD of oxytocin neurons relied on postsynaptic NMDARs. Oxytocin release downstream of NMDAR activation^32^ was not involved, as application of the oxytocin receptor antagonist OTA did not prevent iLTD (**Extended Data Fig. 9d**). NMDAR signaling can lead to a decrease in inhibitory transmission via Ca^2+^-based signal transduction cascades^33, 34^. In addition, NMDARs can alco lead to activation of protein kinase C (PKC)^35^ is involved in dynamin phosphorylation and dynamin-dependent internalization of postsynaptic GABAARs^36, 37^. Indeed, we found that iLTD and change in IPSC/EPSC ratio were blocked by infusing the postsynaptic oxytocin neuron with a membrane-impermeable dynamin inhibitory peptide (i-Dynamin inhibitor) through the recording pipette, but not by a scrambled dynamin inhibitor (i-Dynamin inhibitor control; **Fig. 3h and i, Extended Data Fig. 9e and f**). These findings suggest that PIL opto rapidly triggers iLTD in oxytocin neurons via postsynaptic NMDARs and dynamin signaling, possibly triggering internalization of postsynaptic GABA_A_Rs to decrease inhibitory transmission and increase spiking output.

Increased firing of oxytocin neurons triggered by auditory PIL input in vivo could have substantial impact on maternal behavior relying on the detection of pup calls^17, 22–24, 38^. Specifically, dams with PIL lesions impair pup retrieval behavior^25, 39^. We asked if chemogenetic suppression of PIL input to PVN would affect the behavioral detection by dams of pup calls, as well as the execution of pup retrieval itself. We virally expressed the inhibitory hM4Di receptor, or mCherry as control, in PVN-projecting PIL neurons, injecting wild-type dams with AAVrg-ENN.AAV.hSyn.Cre.WPRE.hGH in PVN and AAV8- hSyn-DIO-hM4D(Gi)-mCherry or AAV8-hSyn-DIO-mCherry in PIL (**Fig. 4a-e**). To specifically test how the PIL-to-PVN pathway regulates the recognition of auditory cues related to pup distress, we tested approach behavior towards speaker playing pup calls in dams expressing hM4Di and administered i.p. with either saline or CNO (**Fig. 4f and g**). Saline-injected dams needed 41.6±5.9 played calls in order to approach the speaker, consistent with our observation that multiple pup calls were required in order to increase the activity of oxytocin cells (**Fig. 4f**); and showed a significant preference for the speaker compared to CNO-injected dams (p=0.002; **Fig. 4g**).

We further tested pup retrieval behavior in dams expressing either hM4Di or mCherry control and administered i.p. with saline or CNO using two different paradigms (**Fig. 4h**). As expected, control dams had significantly lower latencies to approach an isolated crying pup than a speaker playing pup calls (p<0.0001; **Fig. 4i**), which is possibly explained by the use of multiple sensory cues in the pup retrieval test. When hM4Di dams were injected with either saline or CNO and were allowed to remain undisturbed in the nest with their litter before being presented with only one pup at each trial for ten trials total: in average they had a decreased probability to retrieve at later trials without change in the time to retrieval (**Fig. 4h, j, k and l**). As expected, we did not observe this effect when CNO-injected hM4Di dams were presented with five isolated pups at the same time, without being previously in the nest (**Fig. 4h and m**). These results are in line with previous findings that PIL controls pup retrieval behavior^39^ and further indicate that the PIL-to-PVN auditory input is particularly involved in the recognition of infant vocalizations for sustaining maternal performance over time.

We hypothesized that if PIL-to-PVN pathway mediates the persistent activity of oxytocin neurons in dams, then playing pup calls would lead to increased oxytocin release in brain areas involved in pup retrieval and maternal motivation^40, 41^. Downstream oxytocin signaling specifically in the VTA^4, 7, 42, 43^ (**Fig. 5a**) has been shown to control the efficiency of pup retrieval^44^. Indeed, infusion of the OXTR antagonist OTA (0.5 mg/mL) decreased retrieval probability when dams were presented with five isolated pups at the same time (**Extended Data Fig. 10**). This effect is possibly due to a the direct complete inhibition of oxytocin signaling in the VTA which has a more drastic consequence for the efficiency of pup retrieval than a selective chemogenetic inhibition of upstream PIL inputs to PVN, relaying information about pup calls. We therefore hypothesized that PIL inputs to PVN may mediate oxytocin release in the VTA in response to pup calls to ensure reliable pup retrieval behavior. We used a novel genetically-encoded oxytocin sensor, *OXTR-iTango2*^45, 46^ to asses pup calls-triggered oxytocin release in the VTA of dams. This optogenetic system allows for detection of local endogenous oxytocin release, by utilizing viral expression of oxytocin receptors and then optically labeling those neurons after oxytocin receptor signaling (**Fig. 5b**). The *OXTR- iTango2* system requires two synthetic proteins, and their interaction causes the restoration of split TEV protease function. Labelling neuronal populations with *OXTR-iTango2* consists of injecting three viral constructs in an area of interest: AAV1-hSYN-OXTR-TEV-C-P2A-iLiD- tTA (coding for the C-terminus truncated form of the oxytocin receptor, the N-terminus of TEV protease, a blue light-sensitive protein AsLOV2, and tetracycline transactivator protein, tTA), AAV1-hSYN-β-Arrestin2-TEV-N-P2A-TdTomato (coding for β-arrestin2 protein fused with the C-terminus of the split TEV protease and tdTomato), and AAV1-hSYN-TRE-EGFP (coding for tetracycline response element, TRE, and EGFP). The reconstitution of the N- and C-termini of the TEV protease function is ligand-dependent and occurs when oxytocin binds to the oxytocin receptor. Furthermore, TEV cleavage site recognition requires presence of blue light. Thus, for *OXTR-iTango2* labelled neuronal populations to detect endogenous oxytocin release, blue light is delivered via optical fiber inserted in the same area. In the presence of oxytocin in the tissue, paired with blue light illumination, the transactivation domain TRE can trigger EGFP expression. Detection of endogenously-released oxytocin in VTA of female and male mice during social interactions has been previously demonstrated using *OXTR-iTango2*^45^.

We injected wild-type dams unilaterally in the left VTA with *OXTR-iTango2* viral constructs and implanted an optical fiber at the same site (**Fig. 5c**). Dams were exposed to either blue light illumination and pup calls (‘Blue light+pup calls’), blue light and pure tones (‘Blue light+tones’), or to blue light only (‘Blue light only’). Auditory stimuli (pup calls or half-octave pure tones depending on group) were paired with blue light illumination delivered through the optical fiber connected to a blue laser (**Fig. 5d**). No auditory stimuli were presented to dams in Blue light only group. Cells expressing *OXTR-iTango2* constructs in the VTA were labelled in red (tdTomato+) and cells expressing the TRE-EGFP reporter signal were labelled in green (EGFP+), marking neurons which responded to endogenously-released oxytocin (**Fig. 5e**). There were much more tdTomato+/EGFP+ cells in the VTA of Blue light+pup calls compared to Blue light+tones or Blue light only groups (**Fig. 5f-h**). Significantly less tdTomato+/EGFP+ cells were observed when PIL to PVN pathway was inhibited with chemogenetics (**Fig. 5i-k**). This demonstrates that the enhanced firing of PVN oxytocin neurons following pup calls can lead to central oxytocin release in downstream brain areas controlling pup retrieval, such as the VTA, and that this effect is mediated by activation of PIL inputs to PVN.

## Discussion

Here we defined a noncanonical auditory circuit to the hypothalamus, specifically from the PIL to PVN oxytocin neurons, that relays infant distress sounds and is important for auditory-driven maternal behavior. Our data show that this circuit disinhibits oxytocin neurons and enables central oxytocin release, providing a mechanistic understanding of how sensory cues from the offspring can be integrated in maternal endocrine networks to gate hormonal release. We found that specifically oxytocin neurons within the PVN of dams exhibited delayed and long-lasting increase in firing after exposure to pup calls but not pure tones, and that the PIL- to-PVN pathway controls central oxytocin release and efficiency of pup retrieval over time. Persistent activity seems to be hallmark of hypothalamic neurons as they have long integration periods and can be highly interconnected^47–50^. Such stimulus-specific and cell-type-specific sustained activity of oxytocin neurons might serve to maintain maternal arousal and performance over extensive periods, thus ensuring efficient care for the offspring.

We found that parvocellular PVN oxytocin neurons are the main target of PIL inputs. Parvocellular neurons project to the VTA^42^ which is consistent with our results that the PIL to PVN pathway mediates pup call-induced oxytocin release in VTA and that blocking oxytocin signaling in the VTA impairs pup retrieval behavior. Even though parvocellular neurons may be only a small number of the total number of oxytocin cells within the PVN, they innervate an important amount of central areas and have been shown to mediate a wide variety of synaptic mechanisms, sensory processing and behaviors^1, 4, 14, 51–55^. Parvocellular PVN cells have more complex morphology^53^ and potentially innervate multiple brain areas by ramified collaterals^1, 56^. In addition to having a distinct morphology, parvocellular neurons can directly activate magnocellular neurons and have been proposed to operate as “master cells” which control the activity of magnocellular cells and gate oxytocin release^16, 57, 58^. This positions parvocellular neurons to be selectively targeted by sensory inputs, and to subsequently control oxytocin release in different brain areas such as the VTA.

Multiple pup calls were required to activate oxytocin cells and induce speaker approach from dams. This indicates that a series of pup calls act as a natural signal for relieving synaptic inhibition and enabling spiking in oxytocin cells. Hypothalamic inhibition increases during lactation^59^ and might prevent activation of oxytocin neurons by sparce synaptic inputs such as occurring during single pup calls or non-salient auditory stimuli as pure tones. A possible mechanism for the delayed sustained firing of oxytocin cells following pup calls in vivo is postsynaptic NMDAR-dependent decrease in inhibition. NMDARs in oxytocin neurons can be activated in the absence of postsynaptic spiking^60^, suggesting that even though PIL inputs were too weak to induce action potentials, prolonged subthreshold activation of oxytocin cells was sufficient for plasticity induction via NMDARs. Postsynaptic NMDARs also regulate cell excitability^61, 62^ and trigger somatodendritic oxytocin release^60, 63^. In turn, oxytocin signaling controls presynaptic GABA release probability via the postsynaptic release of endocannabinoids and activation of presynaptic CB1 receptor^64^. Therefore, NMDARs can regulate synaptic inhibition in oxytocin cells by both pre- and postsynaptic mechanisms. Transient PIL activation such as occurring during pure tones, however, was unable to promote disinhibition. We further found that pup calls but not pure tones trigger oxytocin release in the VTA via activation of the PIL-to-PVN pathway and that oxytocin signaling in the VTA has been shown controls retrieval behavior. This, in combination with our data, suggests that release of oxytocin in the VTA, via activation of PIL-to-PVN pathway by pup calls, may be involved in sustaining efficient pup retrieval behavior over time. Our findings provide a biological basis for the circuit and synaptic mechanisms that connect sensory cues from the offspring to hormonal release in mothers, perhaps to promote plasticity and amplify neural representations across a broader range of infant cues important for sustaining maternal arousal and performance over time.

## Methods

### Animals

All procedures were approved under NYU School of Medicine IACUC protocols, in accordance with NIH guidelines. Animals were housed in fully-equipped facilities in the NYU School of Medicine Science Building (New York City). The facilities were operated by the NYU Division of Comparative Medicine. All experimental dams (0-3 weeks postpartum) had given birth for the first or second time and were housed with their own or foster pups (P0-P21), or were just weaned (<1 week post-weaning). Surgeries were performed at 0-2 weeks postpartum in dams and >6 weeks of age in virgins. Dams were cohoused with younger pups (P1-5) to maintain maternal behavior after weaning of their own litter. Wild-type C57BL/6N (Taconic, B6-F) dams were used for in vivo and in vitro electrophysiology, anatomy tracings, oxytocin sensor experiments and behavior. Oxytocin:Cre (Jackson, 024234) dams and virgins were used for anatomy tracings and fiber photometry. Oxytocin:Cre x Ai9 dams were used for in vitro electrophysiology and anatomy tracings. Oxytocin:Cre x Ai32 dams were used for in vivo electrophysiology and histology validation. Mice were maintained on a normal 12-h light/dark cycle (dark cycle starts at 6 PM) and given food and water ad libitum. In vivo electrophysiology and behavioral experiments were performed at the end of the light cycle of the animals.

### Stereotaxic surgery

Animals were anesthetized with isoflurane (1–1.5%) and bilaterally injected with 400 nL (in PVN) of either AAV2-EF1a-FLEX-TVA-GFP (Salk Institute Viral Vector Core, Item# 26197, titer: 2.73 × 10^10 vg/mL), EnvA G-Deleted Rabies-mCherry (Salk Institute Viral Vector Core, Item# 32636, titer: 3.78 × 10^7 vg/mL)^65^, AAVrg- ENN.AAV.hSyn.Cre.WPRE.hGH (Addgene, Item# 105553-AAVrg, titer: 7 × 10^12 vg/mL) or AAV5-hSynapsin1-FLEx-axon-GCaMP6s (Addgene, Item# 112010-AAV5, titer: ≥ 7 × 10^12 vg/mL) or AAVDJ-CAG-FLEx- GCaMP6s (Penn Vector Core, Lot# V7340S, titer 3.2 × 10^13); 200 nL (in PIL) of either AAV8-hSyn-DIO-hM4D(Gi)-mCherry (Addgene, Item# 44362-AAV8, titer: 1 × 10^13 vg/mL) or AAV8-hSyn-DIO-mCherry (Addgene, Item# 50459-AAV8, titer: 1 × 10^13 vg/m); 200 nL (in PIL), 600 nL (in AuCx) or 1 µL (in IC) of AAV1-hSyn-hChR2(H134R)-EYFP (Addgene, Item# 26973-AAV1, titer: 1 × 10^13 vg/mL); unilaterally injected in the left hemisphere with 500 nL (in VTA) of *OXTR-iTango2* viral constructs of 1:1:2 ratio: AAV1-hSYN-OXTR-TEV-C-P2A-iLiD-tTA (titer: 2.1 × 10^13 vg/mL), AAV1-hSYN-β-Arrestin2-TEV-N-P2A-TdTomato (titer: 2.57 × 10^14 vg/mL), and AAV1-hSYN-TRE-EGFP (titer: 1.17 × 10^13 vg/mL). Nanoject III (Drummond Scientific, Item# 3-000-207) was used for AuCx, PIL, PVN and VTA injections. Pump 11 Elite Syringe Pump (Harvard Apparatus, Item# HA1100) with a 5 µL Hamilton syringe Model 75 RN SYR (Hamilton Company, Item# 7634-01) was used for IC injections. We waited two weeks after injection of AAV2-EF1a-FLEX-TVA-GFP to inject EnvA G-Deleted Rabies-mCherry in PVN. AAVrg-ENN.AAV.hSyn.Cre.WPRE.hGH (in PVN) and AAV8-hSyn-DIO-hM4D(Gi)-mCherry or AAV8-hSyn-DIO-mCherry (in PIL) were simultaneously injected. We used the following stereotaxic coordinates (in mm): AuCx (-2.54 A-P, +/-4.5 M-L, -0.5 D-V), IC (-5.2 A-P, +/-1.5 M-L, -1.5 D-V), PIL (-2.7 A-P, -1.7 M-L, -3.7 D-V), PVN (-0.72 A-P, +/-0.12 M-L, -4.7 D-V) and VTA (-2.5 A-P, +/-0.4 M-L, -4 D-V).

### In vivo awake cell-attached and tungsten recordings

Oxytocin:Cre x Ai32 or wild-type dams were head-fixed using custom-made 3D-printed headposts and head-fixation frames. A small craniotomy (<1 mm) was performed over the sagittal sinus centered above the left PVN (coordinates in mm: -0.72 A-P, -0.12 M-L) in Oxytocin:Cre x Ai32 dams and above the left PIL (coordinates in mm: -2.7 A-P, -1.7 M-L) in wild-type dams. Cell-attached or whole-cell recordings from optically-identified channelrhodopsin2-expressing oxytocin neurons or other unidentified neurons were obtained from the PVN (4-5.2 mm below the pial surface). Cell-attached recordings using a conventional microelectrode holder or multiunit recordings with tungsten microelectrodes of PIL neurons in wild-type dams were obtained at 3.2-3.7 mm below the pial surface. Pipettes with resistance of 5-6 MΩ and a long taper (∼6 mm) designed for subcortical recordings were made of borosilicate glass capillaries with O.D. 1.5 mm, I.D. 0.86 mm (Sutter Instruments, Item# BF-150-86-10) using P-87 micropipette puller (Sutter Instruments, Item# P-87) and a 3 mm trough filament (Sutter Instruments, Item# FT330B). Patch pipettes contained (in mM):

127 K-gluconate, 8 KCl, 10 phosphocreatine, 10 HEPES, 4 Mg-ATP, 0.3 Na-GTP (osmolality, 285 mOsm; pH 7.2 adjusted with KOH). The pressure of the patch pipette was monitored with manometer (Omega, Item# HHP680). 15-20 mbar pressure was applied when the patch pipette was lowered into the brain and the pressure was adjusted to 1.5-2 mbar when the pipette reached the targeted depth from the pial surface. Cell-attached recordings were obtained with a Multiclamp 700B amplifier (Molecular Devices) and data were acquired with Clampex 10.7 (Molecular Devices), low-pass filtered at 1 kHz, high-pass filtered at 100 Hz and digitized at 20 kHz. Multiunit recordings were obtained using tungsten microelectrodes (Microprobes for Life Science, Item# WE30030.5A5) with 2-3 µm diameter and 0.5 MΩ impedance connected to DAM50 Extracellular Amplifier (World Precision Instruments); data were acquired with Clampex 10.7 (Molecular Devices), low-pass filtered at 3 kHz, high-pass filtered at 300 Hz and digitized at 100 kHz. The tip of the tungsten electrode was coated with DiI (Thermo Fisher Scientific, Item# D282) for histological validation of the recording site.

### In vivo optogenetic identification of PVN neurons

To identify channelrhodopsin2-expressing PVN oxytocin neurons in Oxytocin:Cre x Ai32 dams via channelrhodopsin2-assisted patching during each recording session^66, 67^, we used a Fiberoptic Light Stimulating Device with 465nm blue LED (A-M Systems, Item# 726500) connected to a Fiber Optic Light guide (A-M Systems, Item# 726527). The optic fiber was inserted into the patch pipette via 1.5 mm O.D. Continuous Optopatcher holder (A-M Systems, Item# 663943). Pulses of blue light were delivered through the optic fiber via Digidata 1440A (Molecular Devices) while recording in cell-attached or whole-cell configuration the responses of channelrhodopsin2-positive oxytocin neurons (ChR2+; OT+) or other channelrhodopsin2-negative PVN neurons (ChR2-; OT-). Different steps of light pulses (50 or 200 ms duration) were delivered with increasing intensity from 20 to 100% of full LED power (3 mW at the tip of the fiber). ChR2+ (OT+) neurons responded to light pulses by an increase in their firing rate and spiking probability, while ChR2- (OT-) neurons were not modulated by blue light.

### Fiber photometry

Oxytocin:Cre dams and virgins were bilaterally injected with AAVDJ-CAG-FLEx-GCaMP6s (Penn Vector Core, Lot# V7340S, titer 3.2 × 10^13) in PVN (coordinates in mm: -0.72 A-P, +/-0.12 M-L, -4.7 D-V). Animals were head-posted and a 400 µm optical fiber (Thorlabs, Item# CFMC54L05) was implanted in the left hemisphere slightly above the injection site at -4.5 D-V. Experiments were performed two to three weeks after surgery. Animals were placed in a head-fixation frame within a custom-built soundproof box and photometry was performed with a custom-built rig^17, 68^. A 400 Hz sinusoidal blue light (40-45 µW) from a 470 nm LED (Thorlabs, Item# M470F1) connected to a LED driver (Thorlabs, Item# LEDD1B) was delivered via the optical fiber to excite GCaMP6s. We also used a control 330 Hz light (10 µW) from a 405 nm LED (Thorlabs, Item# M405FP1) connected to a second LED driver.

Light travelled via 405 nm and 469 nm excitation filters via a dichroic mirror to the brain. Emitted light traveled back through the same optical fiber via dichroic mirror and 525 nm emission filter, passed through an adjustable zooming lens (Thorlabs, Item# SM1NR05) and was detected by a femtowatt silicon photoreceiver (Newport, Item# 2151). Recordings were performed using RX8 Multi-I/O Processor (Tucker-Davis Technologies). The envelopes of the signals were extracted in real time using Synapse software (Tucker-Davis Technologies). The analog readout was low-pass filtered at 10 Hz.

### Auditory stimulation

Pup distress vocalizations were recorded from isolated pups aged postnatal day (P) 2-8. Adult vocalizations were recorded during social interactions of male and female mice (> 6 months old). Both pup and adult calls were recorded using an ultrasonic microphone (CM16/CMPA, Avisoft Bioacoustics, Item# 41163, 200 kHz sampling rate; connected to UltraSoundGate 116H recording interface, Avisoft-Bioacoustics, Item# 41163) and de-noised/matched in peak amplitude (Adobe Audition) similar to previous work^34^. To investigate responses to pup calls in individual oxytocin neurons, we monitored baseline firing rates for 1-2 min. For most cells, we then played a set of pup calls consisting of 3-5 individual pup calls (∼1 s duration of each call) repeated for a total of 15-18 calls with a 1 sec delay between calls. Most calls were isolation/distress calls; for one neuron, wriggling calls were played. For the fiber photometry experiments (**Fig. 1k and m, and Extended Data Fig. 3a, b, c, d, i and j)**, we played 4 sets of pup or adult calls (pup calls: 5 calls; adult calls: 3 calls, for a total of 15 calls per set) separated by 1-3 minutes each. For the cells in **Extended Data Fig. 2h-j**, individual calls were played after 1 s baselines, with 3-6 different pup calls were played for 10-15 times each. For studies of pure tones, half-octave (4–64 kHz) or ultrasound half-octave (23-64 kHz) 4 sets of pure tones (10 ms cosine ramp) at 70 dB sound pressure level (SPL) were played for 1 second (**Fig. 1l and n, and Extended Data Fig. 3e, f, g and j**), in a pseudorandom order (repetition rate: 0.2 Hz, total of 15 tones per set, 3-4 sets). Frequency-modulated down-sweeps (“FM sweeps”) which retained key spectrotemporal characteristics of pup vocalizations (72-30 kHz, 200 ms inter syllable interval, 100 ms sweep duration, 0.2 Hz repetition rate, total of 15 sweeps per set, 3-4 sets) were played at 70dB for 4 sets. For the cells in **Extended Data Fig. 3k**, 50 ms pure tones were played. All auditory stimuli were played using RZ6 auditory processor (Tucker-Davis Technologies).

### In vitro whole-cell recordings

Recordings in brain slices were conducted two to three weeks after virus injection. After being anesthetized by isoflurane inhalation, mice were perfused with ice-cold sucrose-based cutting solution containing (in mM): 87 NaCl, 75 sucrose, 2.5 KCl, 1.25 NaH2PO4, 0.5 CaCl2, 7 MgCl2, 25 NaHCO3, 1.3 ascorbic acid, and 10 D-Glucose, bubbled with 95%/5% O_2_/CO_2_ (pH 7.4). The brain was rapidly placed in the same solution and 250 µm slices were prepared with a vibratome (Leica P-1000), placed in warm sucrose-based cutting solution bubbled with 95%/5% O_2_/CO_2_ (pH 7.4), and maintained at 33-35°C for ∼30 min, then cooled to room temperature (22-24°C) for at least one hour before use. For experiments, slices were transferred to the recording chamber and superfused (2.5-3 ml.min^-1^) with artificial cerebrospinal fluid (ACSF, in mM: 124 NaCl, 2.5 KCl, 1.5 MgSO4, 1.25 NaH2PO4, 2.5 CaCl2, 26 NaHCO3 and 10 D-Glucose) at 33°C bubbled with 95%/5% O_2_/CO_2_ (pH 7.4). Neurons were identified with an Olympus 40 x water-immersion objective with TRITC filter. Pipettes with resistance 5-6 MΩ made of borosilicate glass capillaries with O.D. 1.5 mm, I.D.

0.86 mm (Sutter, Item# BF-150-86-10) contained for voltage-clamp recordings (in mM): 130 Cs-methanesulfonate, 1 QX-314, 4 TEA-Cl, 0.5 EGTA, 10 phosphocreatine, 10 HEPES, 4 Mg-ATP, 0.3 Na-GTP (osmolality, 285 mOsm; pH 7.32 adjusted with CsOH), or for current-clamp recordings (in mM): 127 K-gluconate, 8 KCl, 10 phosphocreatine, 10 HEPES, 4 Mg-ATP, 0.3 Na-GTP (osmolality, 285 mOsm; pH 7.2 adjusted with KOH). Somatic whole-cell voltage-clamp and current-clamp recordings were made from PVN oxytocin neurons or PIL neurons with an Multiclamp 200B amplifier (Molecular Devices). Data were acquired with Clampex 10.7 (Molecular Devices), low-pass filtered at 2 kHz and digitized at 20 kHz. To measure synaptic connectivity between areas using channelrhodopsin2-assisted circuit mapping^69^, whole-cell voltage-clamp recordings were made and neurons were held at -70 mV or 0 mV for EPSC or IPCS recordings, respectively. Channelrhodopsin2-expressing axons were activated with 1 ms pulses of full field illumination with 465 nm LED light (Mightex, Item# SLC-AA02-US) repeated at 0.1 Hz. To measure NMDAR-mediated currents, whole-cell voltage-clamp recordings were made in the presence of picrotoxin (50 µM) and neurons were held at +40 mV. Amplitudes of NMDAR-mediated currents was measured 50 ms after optogenetic stimulation.

### Long-term synaptic plasticity experiments

Whole-cell current-clamp or voltage-clamp recordings were made from PVN oxytocin neurons (red fluorescent) of Oxytocin:Cre x Ai9 dams injected with AAV1-hSyn- hChR2(H134R)-EYFP in PIL. Action potentials or synaptic currents were evoked using an extracellular bipolar electrode made of 0.015” silver-chloride filament inserted into a borosilicate theta capillary glass O.D. 1.5 mm, I.D. 1.00 mm (Warner Instruments, Item# TG150-4) filled with ACSF and placed lateral and in close proximity to PVN. To evoke action potentials or postsynaptic potentials in current-clamp mode, or synaptic currents in voltage-clamp mode, electrical pulses (0.1-10 mA and 0.1 ms duration) were delivered at 0.1 Hz with Stimulus Isolator (World Precision Instruments, Item# A365). Repeated optogenetic stimulation of PIL terminals in PVN (PIL opto; 30 Hz during 1 s, repeated at 0.5 Hz for 3 min) was designed to mimic PIL discharge during pup call presentation in vivo. Repeated brief optogenetic stimulation of PIL terminals in PVN (PIL opto short; 30 Hz during 100 ms, repeated at 0.5 Hz for 3 min) was designed to mimic PIL discharge during pure tones presentation in vivo. To examine action potential generation, baselines were recorded for 2-5 min and the intensity of the electrical stimulation was tuned to remain mainly subthreshold, subthreshold activity or evoked action potentials were recorded for 5 or 20 min after PIL opto. For monitoring synaptic currents, neurons were held at -70 mV for EPSCs and 0 mV for IPSCs. Neurons were held at -50 mV during PIL opto. Following PIL opto or PIL opto short, synaptic currents were recorded for 15-20 min. For occlusion experiments, Oxytocin-Cre x Ai9 dams were exposed to playback of pup calls (3-5 individual pup calls, ∼1 s duration of each call, repeated for a total of 15 calls with a 1 sec delay between calls), brains were immediately collected for preparation of PVN slices, and synaptic plasticity recordings were obtained after 1h of slice incubation. For excitability experiments, neurons were recorded in current-clamp configuration and held at ∼-50 mV (close to resting membrane potential). Number of evoked spikes by different amplitudes of intracellular current injection was calculated before and after PIL opto. Recordings were excluded from analysis if the access resistance (Ra) changed >30% compared to baseline. (d(CH[)[¹,Tyr(Me)²,Thr[,Orn[,des- Gly-NH[[)-Vasotocin trifluoroacetate salt (OTA; 1 µM; Bachem, Item# 4031339), DL-2- amino-5-phosphono-pentanoic acid (AP5; 50 µM; Tocris, Item# 0105), 4-Aminopyridine (4- AP; 100 µM; Tocris, Item# 0940), 6,7-Dinitroquinoxaline-2,3-dione (DNQX; 25 µM; Tocris, Item# 0189), (S)-2-Amino-2-methyl-4-phosphonobutanoic acid (MAP4; 250 µM; Tocris, Item# 0711), Tetrodotoxin (TTX; 1 µM; Tocris, Item# 1078) and clozapine N-oxide hydrochloride (CNO; 1 µM; Millipore Sigma, Item# SML2304) were dissolved directly in the extracellular solution and bath applied. Picrotoxin (50 µM; Millipore Sigma, Item# P1675) was dissolved in ethanol and added to the extracellular solution, such that the final concentration of ethanol was 0.1%. Dizocilpine maleate (MK801; 1 mM; Tocris, Item# 0924), Dynamin inhibitory peptide, sequence QVPSRPNRAP (1.5 mM; Tocris, Item# 1774) and Dynamin inhibitory control peptide, sequence PRAPNSRQPV (50 µM; GenScript) were dissolved directly in the intracellular solution.

### Chemogenetic inactivation

Wild-type dams were bilaterally injected with AAVrg-ENN.AAV.hSyn.Cre.WPRE.hGH in PVN and with either AAV8-hSyn-DIO-hM4D(Gi)-mCherry or AAV8-hSyn-DIO-mCherry in PIL. Approach to speaker broadcasting pup calls or pup retrieval behavior were measured at least three weeks after virus injection. 1-2 days before behavior testing, dams were housed in a 26 x 34 x 18 cm cage with pups of P1-5 and nesting material. The day of behavior testing, dams were intraperitoneally injected with 1 mg/kg clozapine N-oxide hydrochloride (CNO) or the equivalent volume of saline and pup retrieval behavior was measured.

For approach to speaker broadcasting pup calls, dams were left in the cage and after verifying that the dam was in the nest with her litter for 1-2 mins, the speaker was turned on and pup calls were played. Speaker was turned off after the dam approached the speaker and a pup was placed in front of the speaker to motivate dams for further trials. For pup retrieval behavior, two to four pups were left in the nest and the remaining pups were kept away from the cage and used for retrieval testing. Ultrasound microphone was used to verify that experimental pups were vocalizing prior to start of the experiment. Retrieval of five pups: dam was placed outside of the cage for 2 min, five pups were placed in the corner opposite to the nest, and the dam was then reintroduced to the cage and placed in the empty nest. The dam was given two trials (2 min per trial) to retrieve all five pups and return them back to the nest. Retrieval of single pup: dam was left undisturbed in the cage and after verifying that the dam was in the nest with her litter for 1-2 mins, one pup was placed in the corner of the cage opposite to the nest. The dam was given ten trials (2 min per trial) to retrieve the displaced pup and return it back to the nest; if the displaced pup was not retrieved within 2 min, the pup was removed and the trial was scored as a failure. Another pup was then placed in an opposite corner and the next trial begun. If dams retrieved pups with 90% accuracy under saline conditions, they were injected with CNO and pup retrieval behavior was tested again after 30-40 min. Behavioral performance (probability to retrieve and time to retrieval) for each dam was compared between saline and CNO conditions. Each session of testing consisted of a baseline set (under saline injection) of 2 trials for five pups retrieval and 10 trials for single pup retrieval, and a post-CNO injection set of 2 or 10 trials, respectively. Experiments were performed under red light.

### Cannula infusion

Wild-type dams were implanted with an infusion cannula (26 GA, 5 mm, Plastics One, Item# C315GAS-5/SPC) over the VTA. The day of the behavioral testing, 1 µL of the OXTR antagonist OTA (0.5 mg/mL) or equal volume of saline was infused over 2 min with Pump 11 Elite Syringe Pump (Harvard Apparatus, Item# HA1100). Pup retrieval test started 5 min following infusion. Behavioral testing was performed in different days for OTA and saline.

### Oxytocin sensor

Wild-type dams were unilaterally injected in the left hemisphere with *OXTR-iTango2*^1, 2^ viral constructs (1:1:2 ratio; AAV1-hSYN-OXTR-TEV-C-P2A-iLiD-tTA, AAV1-hSYN-β- Arrestin2-TEV-N-P2A-TdTomato, and AAV1-hSYN-TRE-EGFP) in VTA (coordinates in mm: -2.5 A-P, -0.4 M-L, -4 D-V). A 200 µm optical fiber (Thorlabs, Item# CFMXB05) was also implanted in the VTA using the same coordinates and a head-post was installed. In a subset of experiments, dams were in addition bilaterally injected with AAVrg-ENN.AAV.hSyn.Cre.WPRE.hGH in PVN and with AAV8-hSyn-DIO-hM4D(Gi)-mCherry in PIL. Pup calls, pure tones, and blue light exposure was performed three weeks after surgery. Dams were head-fixed during experiments. Animals in Blue light + pup calls group were exposed to pup calls (5 different pup calls; ∼1 s duration; 0.5 Hz repetition rate; RZ6 auditory processor, Tucker-Davis Technologies) and paired with blue light illumination (5 s ON/15 s OFF; DPSS Blue 473 nm laser, Opto Engine, Item# MBL-F-473) for 45 min. Animals in Blue light + tones group were exposed to half-octave pure tones (ranging 4–64 kHz; 1 s duration, 10 ms cosine ramp, 70 dB SPL, 0.2 Hz repetition rate) played in a pseudorandom order with blue light illumination (5 s ON/15 s OFF) for 45 min. Animals in Blue light only group were exposed to blue light illumination only (5 s ON/15 s OFF) for 45 min. Animals in Blue light + pup calls + h4DM(Gi) + CNO group were injected with 1 mg/kg CNO 30 min prior to exposure to pup calls.

### Histology and imaging

Animals were deeply anaesthetized with isoflurane inhalation followed by intraperitoneal injection (0.1 ml per 10 g) of a ketamine–xylazine mixture containing 15 mg/ml ketamine and 5 mg/ml xylazine, and transcardially perfused with phosphate buffered saline (PBS) followed by 4% paraformaldehyde (PFA) in PBS. Brains were immersed overnight in 4% PFA followed by immersion in 30% sucrose for two nights. Brains were embedded with Tissue-Plus™ O.C.T. Compound medium (Thermo Fisher Scientific, Item# 23-730) and sectioned at 50 µm thickness using a cryostat (Leica). After cryosectioning, brain sections were washed with PBS (3 x 10 min at room temperature) in a staining jar and incubated for 2 h at room temperature in blocking solution containing 5% normal goat serum (Millipore Sigma, Item# G6767) in 1% Triton X-100 (Millipore Sigma, Item# 11332481001) dissolved in PBS, followed by 48 h incubation at room temperature with primary antibodies at 1:1000 or 1:500 dilution in 3% normal goat serum in 1% Triton X-100 dissolved in PBS. Sections were washed with PBS (3 x 10 min at room temperature) and incubated for 2 h at room temperature in Alexa-Fluor-conjugated secondary antibodies diluted at 1:500 in PBS. Following washing with PBS (2 x 10 min at room temperature), sections were incubated with Hoechst 33342 (Thermo Fisher Scientific, Item# H3570) at 1:1000 for nuclear staining. After washing with PBS (2 x 10 min at room temperature), slides were coverslipped using Fluoromount-G (SouthernBiotech). Slides were examined and imaged using a Carl Zeiss LSM 700 confocal microscope with four solid-state lasers (405/444, 488, 555, 639 nm) and appropriate filter sets. Primary antibodies: GFP Tag Polyclonal Antibody (Thermo Fisher Scientific, Item# A10262), Anti-mCherry antibody (Abcam, Item# ab167453), Anti-Oxytocin Antibody (Millipore Sigma, Item# AB911). Secondary antibodies: Goat anti-Rabbit IgG (H+L) Cross-Adsorbed Secondary Antibody, Alexa Fluor 555 (Thermo Fisher Scientific, Item# A-21428), Goat anti-Chicken IgY (H+L) Secondary Antibody, Alexa Fluor 488 (Thermo Fisher Scientific, Item# A-11039).

### Electrophysiology

Off-line analysis was performed with Clampfit 10.7 (Molecular Devices). Spikes were automatically detected by threshold crossing. To investigate long-term responses to pup calls, normalized firing rate was computed by calculating percentage change in firing rate each minute. To investigate responses to individual pup calls, firing rate was measured throughout the call duration plus 300-400 ms, compared to spontaneous firing 1 s before call onset. To investigate responses to pure tones, firing rate was measured 100 ms from onset of the tone compared to spontaneous rate 100 ms before tone onset. Z-scores were computed by calculating the call-evoked firing rate relative to the spontaneous firing rate: z = (µ_evoked_ –µspontaneous) / σspontaneous^22^.

### Fiber photometry

Data from both the 470 (signal) and 405 (control) wavelength channels were independently filtered using a sliding window^70^. To normalize the scale of the control channel, a least- squares regression of the 470 and 405 wavelength channels was performed. Baseline corrections were estimated by subtracting the normalized control channel from the signal channel. The calculation of ΔF/F was carried out using the following equation: (F-F_0_)/F_0_, where F_0_ represents the baseline signal detected by a first order polynomial fitting. The analysis of all animals included the ’Pre’ phase, which was defined as the 20 seconds prior to the onset of pup calls, the ’Calls’ phase, which corresponded to the period of call playback, and the ’Post’ phase, which covered the 20-40 seconds after call offset.

### Oxytocin sensor

The *OXTR-iTango2*-labeled population in the VTA was categorized into four types based on red and green fluorescent signals from *post hoc* confocal imaging data analysis. Individual regions of interest (ROIs) for cells were semi-automatically drawn using a custom algorithm (ImageJ) based on fluorescence intensity, cell size, and cell shape. The average red (R) and green (G) fluorescent signals were calculated for each ROI and were divided by the mean background value (R_0_ and G_0_ for red and green channels, respectively) outside of the ROIs for normalization. ROIs were allocated into four different quadrants divided by thresholds in two fluorescent colors (x-axis: red tdTomato signal, red threshold value: -1.5; y-axis: green EGFP TRE reporter signal, green threshold value: 0.585). ROIs with red signals above or below the red threshold value were categorized as tdTomato+ or tdTomato-, respectively. ROIs with green signals above or below the green threshold value were categorized as EGFP+ or EGFP-, respectively.

### Statistics

Electrophysiology data analysis was performed using Clampfit (Molecular Devices). Fiber photometry analysis was performed using MATLAB 2017b (MathWorks). Image analysis was conducted using NIH ImageJ. We used Wilcoxon matched-pairs signed rank two-tailed test in **Fig. 1m, 1n, 3c, 3d, 3g and 4d, and EDF 1f, 2g, 2i, 3a, 3b, 3c, 3d, 3e, 3f, 3g, 3h, 4d, 5d, 5e, 7f, 7g, 7h and 7i**. Friedman test was used in **Fig. 1g**. One-sample two-tailed Student’s t-test was used in **Fig. 1h, 3e, 3f and 4e, and EDF 4e, 6e, 8b, 8c, 8e, 8f, 8h, 8i, 9a, 9b, 9c, 9d, 9e and 9f**. Mann-Whitney two-tailed test was used in **Fig. 3e, 3f, 3h, 4i, 4l and 4m, and EDF 1g, 4f, 8d, 8g, 9a, 9b, 9c, 9d, 9e and 9f**. One-way ANOVA was used in **Fig. 5k, and EDF 1b, 1d, 1e and 2j**. Fisher’s two-tailed test was used in **Fig. 4g and 4k** and in **EDF 10c**. Two-tailed Binomial test was used in **Fig. 4g.** All sample sizes and definitions as well precision measures (mean, SEM or SD) are provided in figure legends. Statistical tests and graph generation were performed using Prism 9 (GraphPad) or MATLAB 2017b (MathWorks).

### Data and code availability

Information about resources, reagents used, and requests for code should be directed to and will be fulfilled by the Lead Contacts, Silvana Valtcheva and Robert C. Froemke.

## Acknowledgements

We thank I. Carcea, E. Glennon, K. V. Kuchibhotla, J. K. Schiavo and S. C. Song for comments, discussions and technical assistance. We thank the NYU Langone Microscopy Core for experimental and technical support. We thank D. Rinberg for sharing the custom-made 3D-printed headpost and head-fixation frame design. We thank D. Lin for help with the code for analysis of the fiber photometry data. Initial aliquots of the EnvA G-Deleted Rabies-mCherry (SADΔG-mCherry) and helper AAV2-EF1a-FLEX-TVA-GFP viruses were a gift from G. Fishell. Illustrations in Fig. 1a and j, Fig. 5d and i, Extended Data Fig. 4a, Extended Data Fig. 6a and Extended Data Fig. 10b were made by Shari E. Ross.

## Author information

### Affiliations

Skirball Institute for Biomolecular Medicine, New York University School of Medicine, New York, NY 10016, USA.

Silvana Valtcheva, Habon A. Issa, Chloe J. Bair-Marshall, Kathleen A. Martin & Robert C. Froemke

Neuroscience Institute, New York University School of Medicine, New York, NY 10016 USA.

Silvana Valtcheva, Habon A. Issa, Chloe J. Bair-Marshall, Kathleen A. Martin, Yiyao Zhang & Robert C. Froemke

Department of Otolaryngology, New York University School of Medicine, New York, NY 10016, USA.

Silvana Valtcheva, Habon A. Issa, Chloe J. Bair-Marshall, Kathleen A. Martin & Robert C. Froemke

Department of Neuroscience and Physiology, New York University School of Medicine, New York, NY 10016, USA.

Silvana Valtcheva, Habon A. Issa, Chloe J. Bair-Marshall, Kathleen A. Martin & Robert C. Froemke

Center for Neural Science, New York University, New York, NY 10003, USA.

Silvana Valtcheva, Habon A. Issa, Chloe J. Bair-Marshall, Kathleen A. Martin & Robert C. Froemke

Solomon H. Snyder Department of Neuroscience, Johns Hopkins University School of Medicine, Baltimore, MD 21205, USA.

Kanghoon Jung & Hyung-Bae Kwon

### Funding

This work was funded by a Fyssen Foundation Postdoctoral Fellowship, Leon Levy Foundation Postdoctoral Fellowship and Brain & Behavior Research Foundation NARSAD Young Investigator Award (S.V.); a T32 MH019524 Training in Systems and Integrative Neuroscience (H.A.I.); a Natural Sciences and Engineering Research Council of Canada PGS-D fellowship (C.J.B.-M.); an NSF Graduate Research Fellowship (K.A.M.); DP1MH119428 (H.-B.K.); the BRAIN Initiative (NS107616, Y.Z. and R.C.F.); Program Projects Grant (NS074972), NICHD (HD088411), NIDCD (DC12557), and a Howard Hughes Medical Institute Faculty Scholarship (R.C.F.).

### Contributions

S.V. performed in vivo cell-attached, whole-cell and tungsten recordings, fiber photometry, in vitro whole-cell recordings, behavior for chemogenetic inactivation studies, oxytocin sensor experiments, viral injections, histology, image acquisition and data analysis of electrophysiology and behavior experiments. H.A.I. performed fiber photometry, behavior for chemogenetic inactivation and cannula infusion studies. C.J.B.-M. performed fiber photometry and in vitro whole-cell recordings for the scrambled dynamin inhibitor experiments. H.A.I., K.A.M. and Y.Z. wrote code and performed analysis of the photometry recordings. K.J. and H.-B.K. contributed with design of viral constructs and data analysis for the oxytocin sensor. S.V. and R.C.F. designed the study and wrote the paper.

### Corresponding authors

Correspondence to Silvana Valtcheva and Robert C. Froemke.

### Ethics declarations

#### Competing interests

The authors declare no competing interests.

## Extended data figures and tables

**Extended Data Figure 1.**
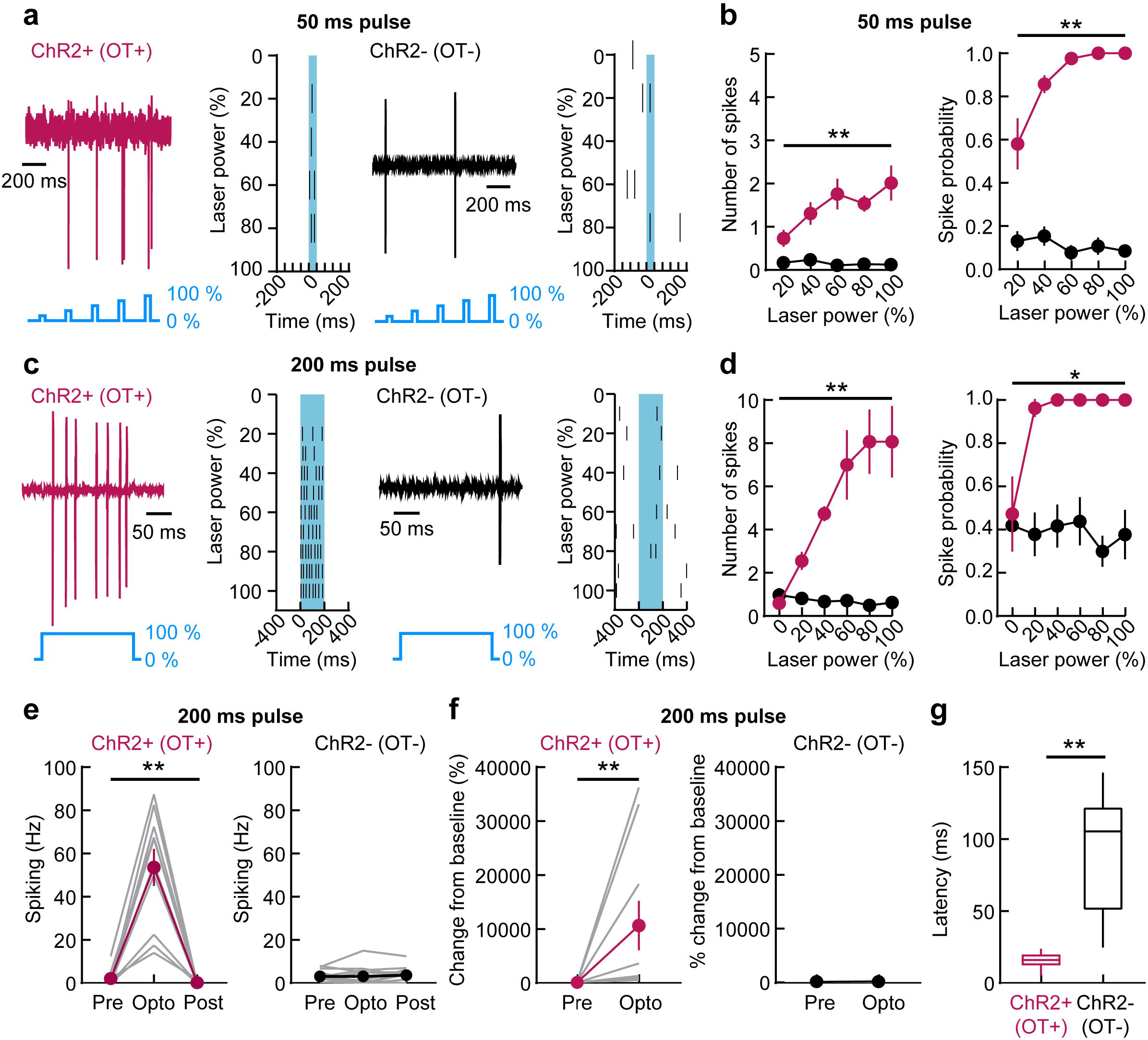
Identification of ChR2+ (OT+) and ChR2- (OT-) neurons. (a) Sample traces and raster plots of cell-attached recordings from one ChR2+ (OT+, left) and one ChR2- (OT-, right) neuron showing reliable activation of ChR2+ (OT+) neurons in response to 50 ms pulses of blue light at 5 Hz (20-100% laser power). (b) Increase in the number of spikes (left) of ChR2+ (OT+; pink circles; n=7 neurons, N=5 dams, p=0.002, one-way ANOVA) and spike probability (right; p=0.009) in response to 50 ms light pulse steps. Number of spikes of ChR2- (OT-; black circles; n=13, N=3, p=0.11) neurons, as well as spike probability (p=0.13) was not modulated. (c) Sample traces and raster plots of cell-attached recordings of one ChR2+ (OT+, left) and one ChR2- (OT-, right). ChR2+ (OT+) neuron was reliably activated in response to 200 ms pulses of blue light (0-100% laser power). (d) Increase in the number of spikes (left) of ChR2+ (OT+; pink circles; n=9, N=6, p=0.002, one-way ANOVA) and spike probability (right; p=0.01) in response to 200 ms light pulse steps. Number of spikes of ChR2- (OT-; black circles; n=13, N=3, p=0.49) neurons, as well as spike probability (p=0.62) was not modulated. (e) Increase in firing rate of ChR2+ (OT+; n=10, N=5, p=0.0001, one-way ANOVA) but not of ChR2- (OT-; n=13, N=3, p=0.68) neurons in response to 200 ms light pulse of 100% laser power (‘Opto’) compared to their baseline firing rate immediately preceding (‘Pre’) and immediately after (‘Post’) the light pulse. (f) Change of firing rate of ChR2+ (OT+; n=10, N=5, p=0.002, Wilcoxon matched-pairs signed rank two-tailed test) but not of ChR2- (OT-; n=13, N=3, p=0.91) neurons in response to 200 ms light pulse of 100% laser power (‘Opto’) compared to baseline firing immediately preceding the light pulse (‘Pre’). (g) Latency to first spike was significantly shorter in ChR2+ (OT+; n=14, N=6, p<0.0001, Mann-Whitney two-tailed test) compared to ChR2- (OT-; n=8, N=2) neurons in response to 200 ms light pulse of 100% laser power. ‘First spike’ in ChR2- (OT-) cells was not light- evoked but occurred spontaneously. Data reported as mean±SEM. *p<0.05. **, p<0.01.

**Extended Data Figure 2.**
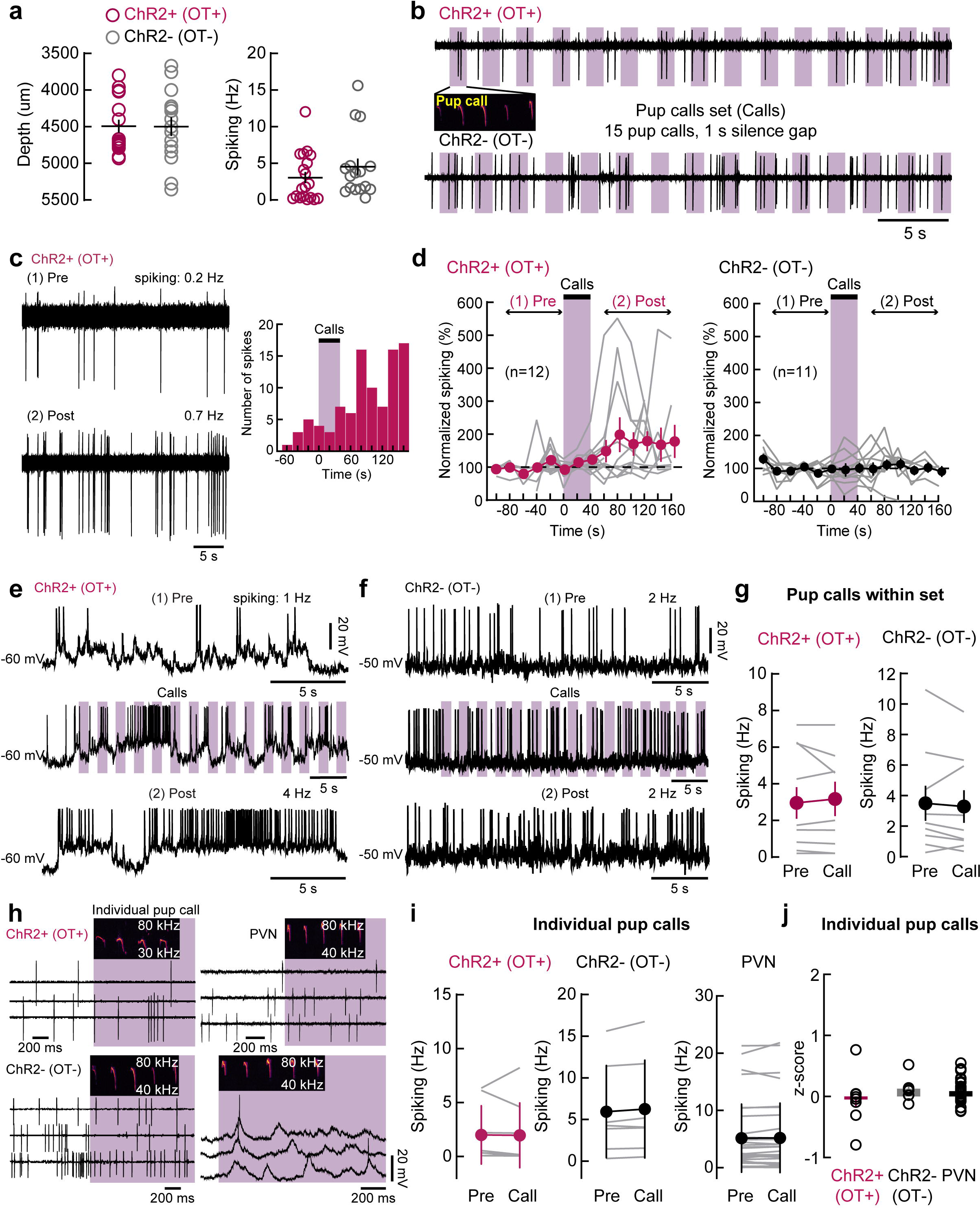
In vivo responses to individual pup calls. (a) Location and firing rate of cell-attached (n=19 neurons, N=8 dams) and whole-cell (n=1) recordings of ChR2+ OT+ neurons and ChR2- neurons (OT-, cell-attached: n=16, N=8; whole-cell: n=2, N=2). (b) Sample traces of whole-cell recordings of one ChR2+ (OT+; upper trace) and one ChR2- (OT-; lower trace) neuron during call playback (‘Calls’). Pink bars, individual pup calls. (c) Left, Cell-attached recording of one ChR2+ (OT+) neuron before pup call onset (1, ‘Pre’), and around 2 min after call onset (2, ‘Post’). Right, Peristimulus time histogram. Bins: 20 s. (d) Timeline of ChR2+ (OT+) and ChR2- (OT-) responses. Period before (1, ‘Pre’) and after (2, ‘Post’) pup call onset. (**e, f**) Sample traces of whole-cell recordings of one ChR2+ (OT+; **e**) and one ChR2- (OT-; **f**) neuron showing baseline spiking activity preceding onset of pup calls (1, ‘Pre’, upper trace), activity during playback of a set of pup calls (‘Calls’, middle trace) and activity after pup calls playback (2, ‘Post’, lower trace). Note increased firing rate for ChR2+ (OT+) neuron but not ChR2- (OT-) neuron. (g) ChR2+ (OT+) did not respond to individual pup calls within a set. ‘Pre’, average spiking rate of cell-attached recordings during all baseline periods immediately preceding each call within the set. ‘Call’, average spiking rate during pup call stimulus for each call within the set. Neither ChR2+ (OT+; n=9, N=6, p=0.38, Wilcoxon matched-pairs signed rank two-tailed test), nor ChR2- (OT-; n=9, N=5, p=0.30) neurons increased their firing rate during individual calls (‘Call’) compared to baseline (‘Pre’). (**h-j**) ChR2+ (OT+) did not respond to presentation of individual pup calls on a trial-by-trial basis. (**h**) Sample traces of cell-attached recordings of one ChR2+ (OT+), one ChR2- (OT-) and one unidentified PVN neuron, as well as whole-cell recording of a PVN neuron during trial-by-trial individual pup call presentation. (**i**) No increase in the firing rates of either ChR2+ (OT+; n=8, N=2, p=0.38, Wilcoxon), ChR2- (OT-; n=7, N=3, p=0.08), or unidentified PVN (cell-attached: n=26, N=13; whole-cell: n=6, N=4, p=0.48) neurons during individual pup calls (‘Call’) compared to baseline (‘Pre’) on a trial-by-trial basis. (**j**) No difference in the z-scores of spiking responses of cell-attached and whole-cell recordings during individual pup calls in ChR2+ (OT+; n=8, N=2, p=0.30, one-way ANOVA), ChR2- (OT-; n=7, N=4), and unidentified PVN (cell-attached: n=21, N=11; whole-cell: n=6, N=4) neurons. Data reported as mean±SEM. *p<0.05.

**Extended Data Figure 3.**
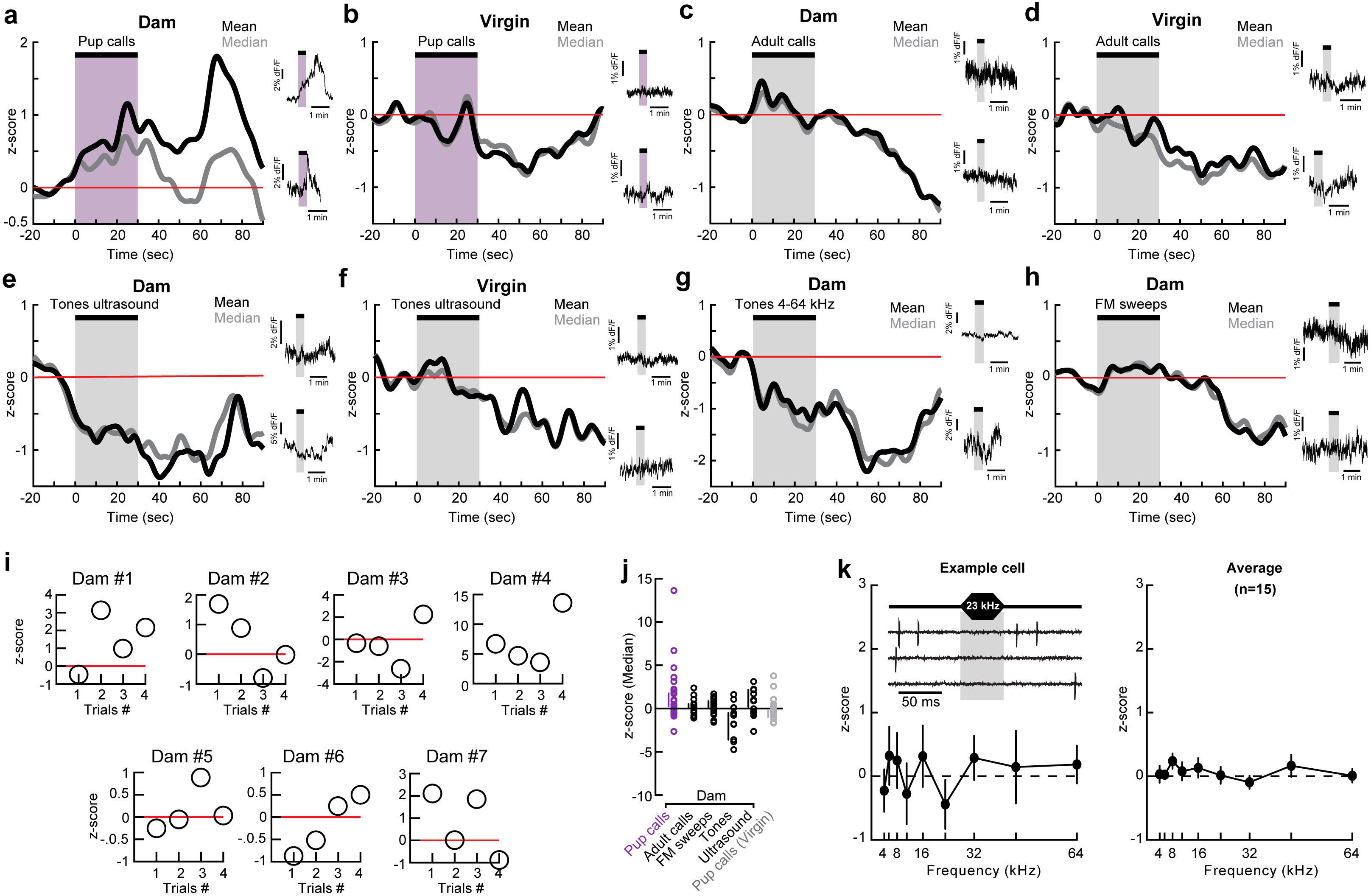
In vivo responses to auditory stimuli. (**a-h**) Z-scores of fluorescence activity during fiber photometry recordings of dams and virgins in response to auditory stimuli. Pup calls trigger sustained increase in the activity of oxytocin neurons in dams (**a**; N=7; Pre vs Post: p=0.0179; Wilcoxon matched-pairs signed rank one-tailed test) but not virgins (**b**; N=4; p=0.4954; Wilcoxon). No responses to adult calls (**c**, N=4 dams; p=0.5966; and **d**, N=3 virgins; p=0.0923; Wilcoxon) or ultrasound pure tones (**e**, N=3 dams; p=0.3776; and **f**, N=3 virgins; p=0.4697; Wilcoxon). No response to pure tones (**g**; N=3; p=0.1099; Wilcoxon) or FM sweeps (**h**; N=5; p=0.1956; Wilcoxon) in dams. Insets represent fluorescence activity during a single trial from two different animals per condition. (i) Responses to pup calls across dams and trials. (j) Average z-score responses to auditory stimuli for individual trials. (k) PVN neurons did not respond to individual pure tones: (**Left**) Example cell-attached recording of one PVN neuron in response to 23 kHz tone presentation and tuning profile of pure tone frequency responses in this cell; (**Right**) Average tuning profile of pure tone frequency responses in PVN cells (n=15 neurons, N=9). Data reported as median±95% CI (j) or as median or mean±SEM (k). *p<0.05.

**Extended Data Figure 4.**
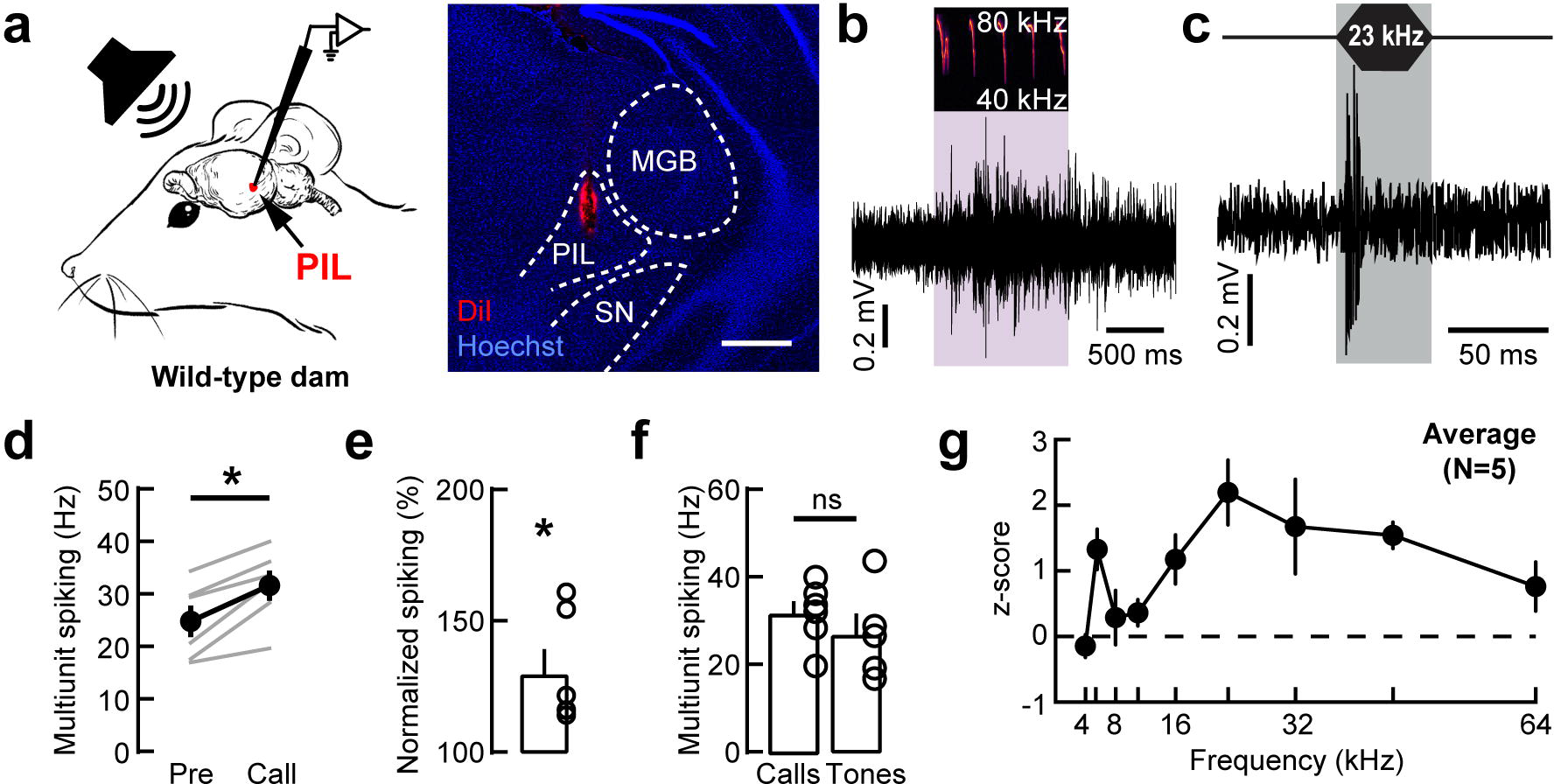
Auditory responses in PIL. (a) Left, experimental setup showing in vivo multiunit recordings via tungsten electrode in PIL of awake head-fixed wild-type dams while playing pup calls from an ultrasound speaker. Right, validation of PIL recording site by coating tungsten electrode tip with DiI. Scale, 500 µm. MGB, medial geniculate body of the thalamus; PIL, posterior intralaminar nucleus of the thalamus; SN, substantia nigra. (**b-g**) In vivo activation of PIL during playback of pup calls and pure tones. Sample trace of stimulus-evoked PIL multiunit spiking activity during individual pup call (**b**) and 23 kHz tone (**c**) presentation. Note the increase in PIL activity during the entire duration of the pup call (>1 s) compared to transient activation during pure tones. Pup calls increased the firing frequency of PIL neurons (**d**; N=6 dams, p=0.03, Wilcoxon matched-pairs signed rank two-tailed test) which corresponded to a significant increase from baseline values (**e**; p=0.02, one-sample two-tailed Student’s t-test). (**f**) There was no difference in the frequency of multiunit spiking during pup calls and pure tones playback (p=0.33, Mann-Whitney two-tailed test). (**g**) Tuning profile of pure tone frequency responses in PIL (N=5). Data reported as mean±SEM. *p<0.05. ns: not significant.

**Extended Data Figure 5.**
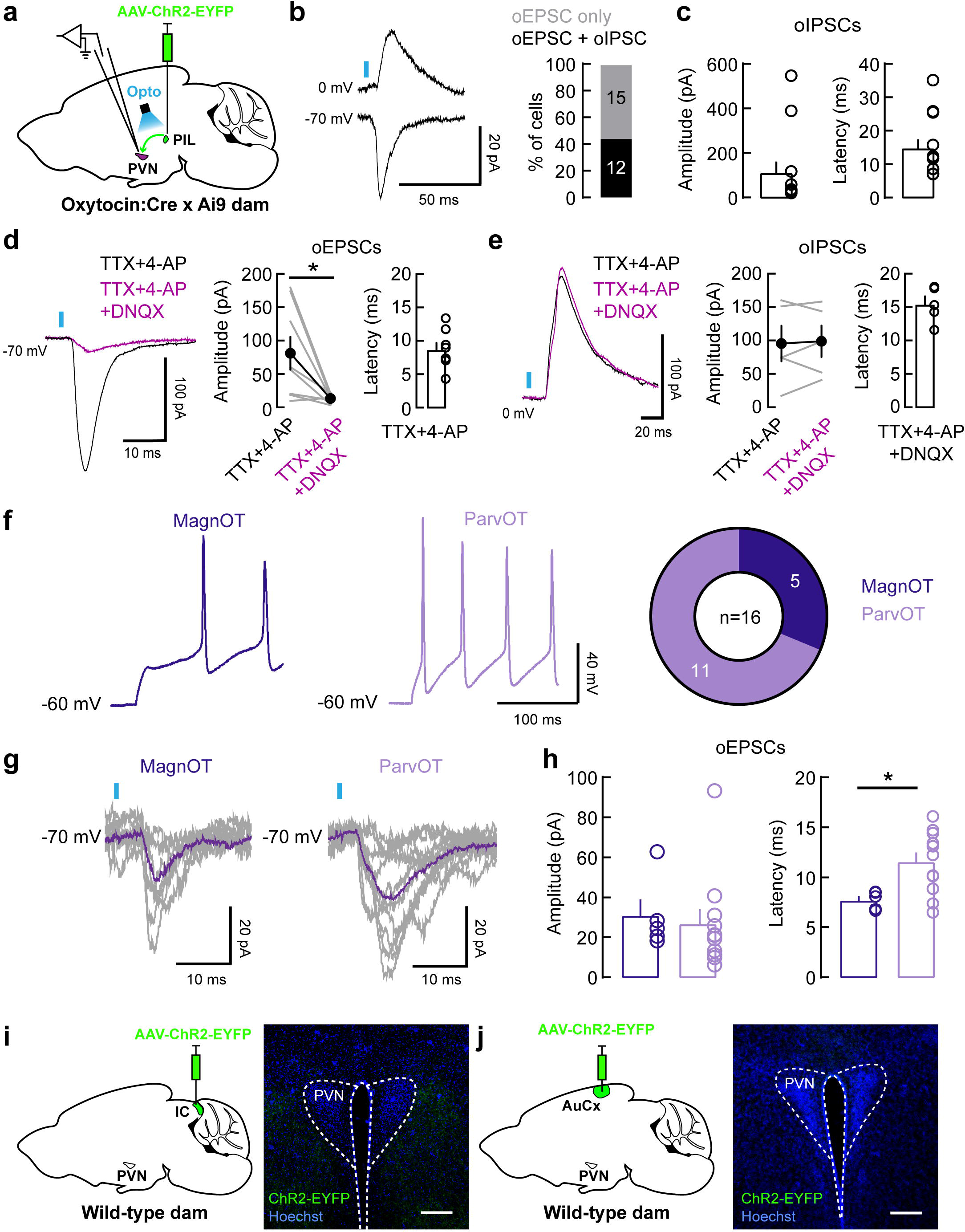
Auditory projections to PVN. (a) Schematic showing injection of AAV1-hSyn-hChR2(H134R)-EYFP in PIL of Oxytocin:Cre x Ai9 dams prior to whole-cell recordings from oxytocin neurons (tdTomato+) in PVN brain slices. PIL, posterior intralaminar nucleus of the thalamus; PVN, paraventricular nucleus of the hypothalamus. (**b and c**) PVN oxytocin neurons receive mainly glutamatergic input from the PIL. Percentage of optogenetically-evoked excitatory (oEPSCs) and inhibitory (oIPSCs) currents in oxytocin neurons triggered by optogenetic stimulation of PIL axons (**b**) and characterization of oIPSCs (**c**). (**d and e**) PIL inputs to oxytocin cells are monosynaptic. (**d**) Example traces and summary graph showing oEPSCs in the presence of TTX and 4-AP and their inhibition by DNQX (n=8, p=0.0156, Wilcoxon matched-pairs signed rank test). (**e**) Example traces and summary graph showing oIPSCs in the presence of TTX and 4-AP; DNQX had no effect on oIPSCs amplitude (n=5, p=0.6250, Wilcoxon). (**f-h**) Parvocellular PVN oxytocin neurons are the main target of input from the PIL. Magnocellular (MagnOT) and parvocellular (ParvOT) oxytocin neurons were characterized by their signature spiking patterns in current-clamp mode (**f**; left). 5/16 MagnOT and 11/16 ParvOT cells received inputs from the PIL (**f**; right). (**g and h**) Characterization of oEPSCs in MagnOT and ParvOT neurons triggered by optogenetic stimulation of PIL axons. There was no difference in oEPSCs amplitude (**h**, left; p=0.51) but the latency of oEPSCs in ParvOT cells was longer (**h**, right; p=0.03). (g) PVN does not receive input from IC. Left, injection of AAV1-hSyn-hChR2(H134R)- EYFP in IC of wild-type dams. Right, no EYFP staining was found in PVN, suggesting that IC does not project to PVN. Scale, 200 µm. N=3. IC, inferior colliculus. (h) PVN does not receive input from AuCx. Left, injection of AAV1-hSyn-hChR2(H134R)- EYFP in AuCx of wild-type dams. Right, no EYFP staining was found in PVN, suggesting that AuCx does not project to PVN. Scale, 200 µm. N=2. AuCx, auditory cortex. Data reported as mean±SEM. *p<0.05.

**Extended Data Figure 6.**
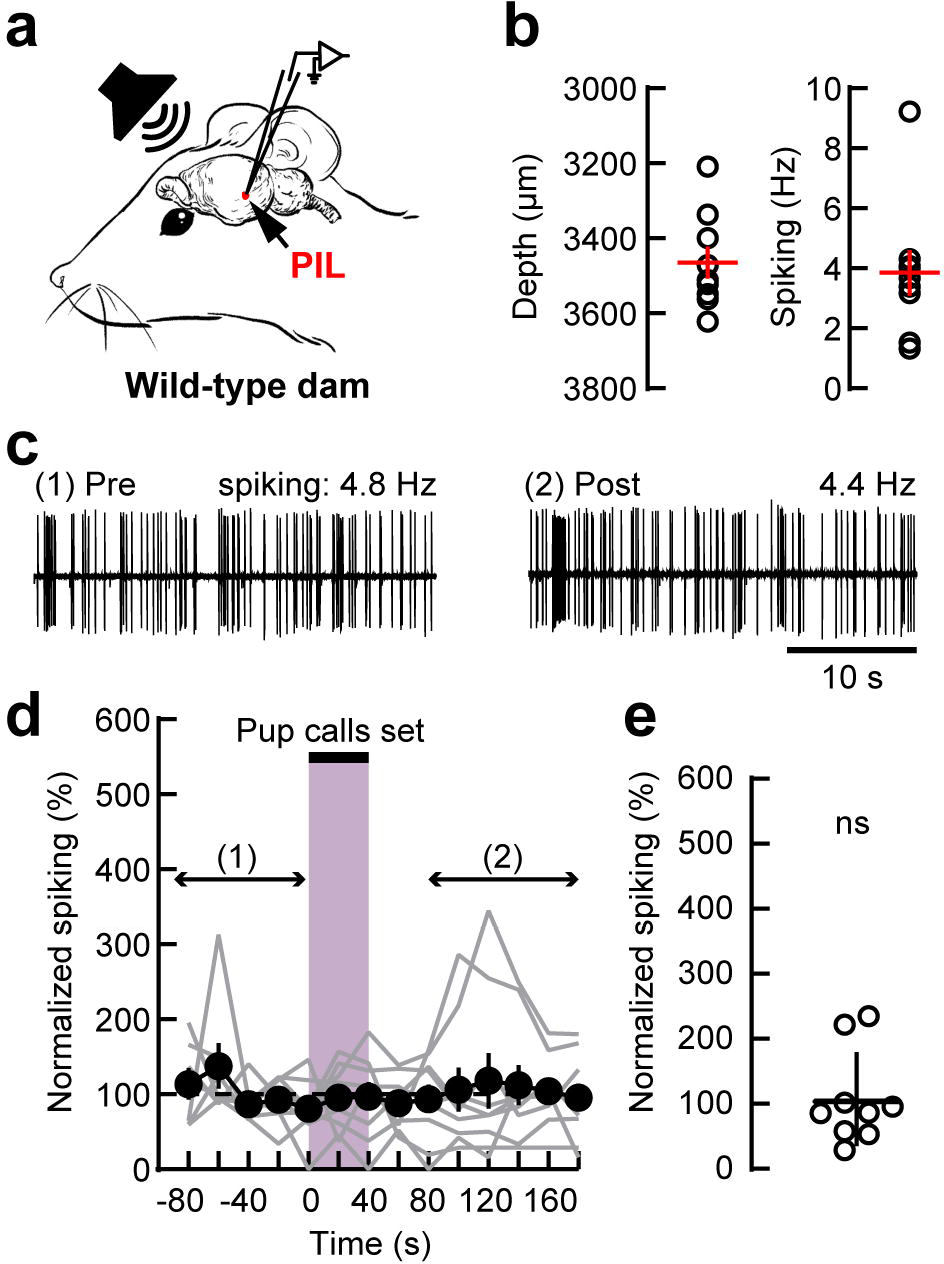
PIL neurons do not exhibit sustained increase in firing following pup calls in vivo. (a) Experimental setup showing in vivo cell-attached recordings in PIL of awake wild-type dams while playing pup calls from an ultrasound speaker. PIL, posterior intralaminar nucleus of the thalamus. (b) Location (depth from pia) and firing rate of PIL neurons (n=9 neurons; N=4 dams). (**c-e**) PIL neurons did not modulate their firing rate following playback of a set of pup calls (15 pup calls, 1 second gap in between calls). (**C**) Sample traces from a cell-attached recording of one PIL neuron showing its baseline firing rate immediately preceding (1, ‘Pre’) and at 90 sec after the onset of pup calls playback (2, ‘Post’). Firing rates during baseline and after pup calls were calculated over 1-2 min. (**d, e**) PIL neurons did not exhibit persistent increase in baseline firing following pup calls, as calculated between 80-160 sec after onset of pup call playback (**e**, n=9, N=4, p=0.77, one-sample two-tailed Student’s t-test). Data reported as mean±SEM. ns: not significant.

**Extended Data Figure 7.**
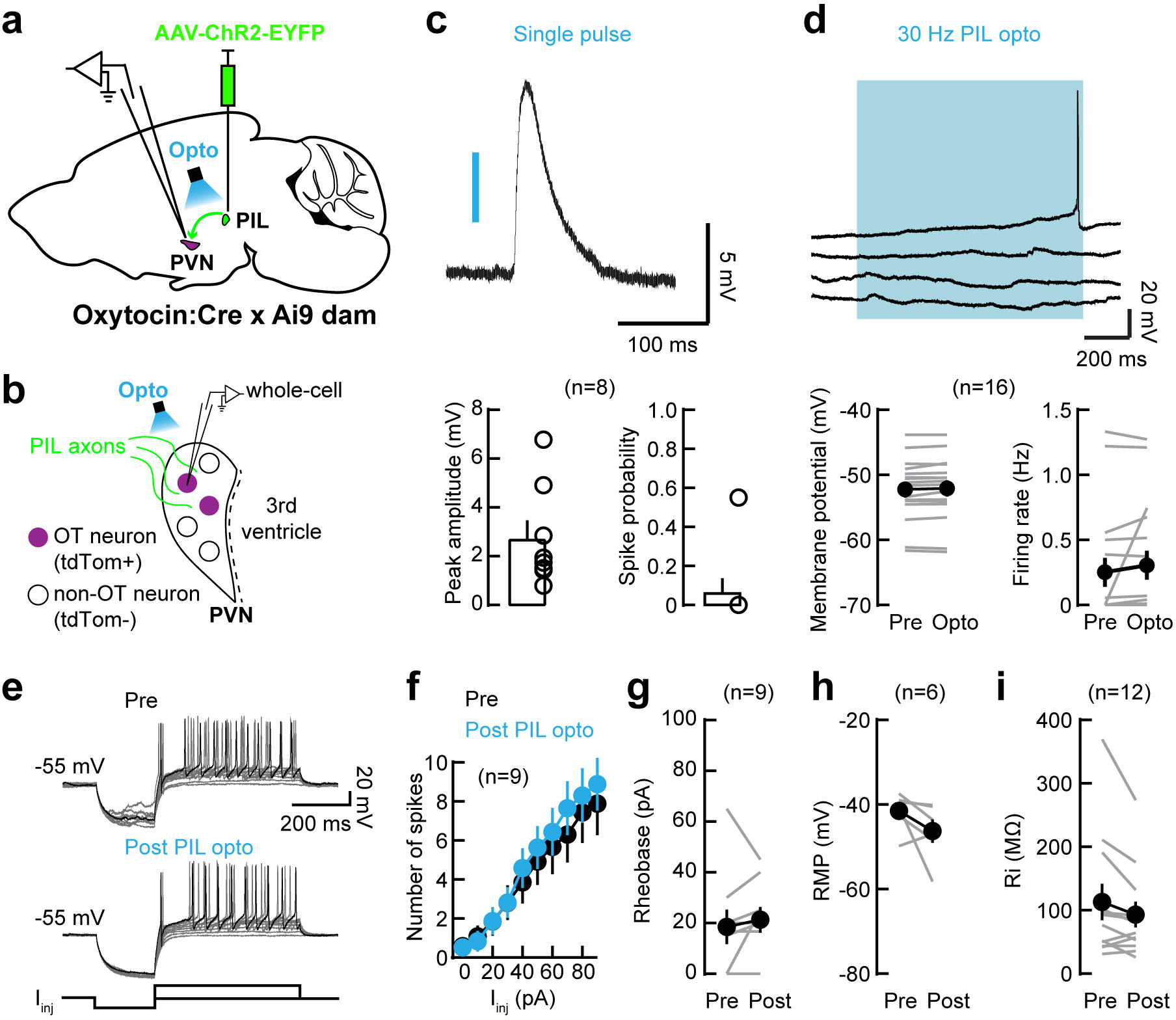
PIL inputs to PVN do not induce postsynaptic spiking or affect excitability of oxytocin neurons. (a) Schematic showing injection of AAV1-hSyn-hChR2(H134R)-EYFP in PIL of Oxytocin:Cre x Ai9 dams prior to whole-cell recordings from oxytocin neurons (tdTomato+) in PVN brain slices. PIL, posterior intralaminar nucleus of the thalamus; PVN, paraventricular nucleus of the hypothalamus. (b) Whole-cell recordings from tdTomato+ oxytocin neurons in PVN slices, optogenetic stimulation of PIL axons and placement of the extracellular stimulation electrode. (**c and d**) Optogenetic stimulation of PIL axons in PVN does not induce postsynaptic spiking in oxytocin neurons. (**c**) Single pulse of optogenetic stimulation triggered postsynaptic potentials but did not induce spiking in oxytocin cells. (**d**) Repeated optogenetic stimulation of PIL axons (‘PIL opto’) did not trigger depolarization of oxytocin cells (p=0.10) and did not induce postsynaptic spiking (p=0.26). (**e, f**) No change in the number of spikes in oxytocin neurons in response to 20 pA steps of intracellular current injection before (‘Pre’) or after (‘Post’) PIL opto. Sample traces (**e**) and summary (**f**; n=9 neurons, p>0.44, Wilcoxon matched-pairs signed rank two-tailed test). (**g-i**) No change in the intrinsic properties of oxytocin neurons after PIL opto, in terms of rheobase (**g**; n=9; p=0.38, Wilcoxon), resting membrane potential (**h**; n=6; p=0.16, Wilcoxon), or input resistance (**i**; n=12; p=0.06, Wilcoxon). Data reported as mean±SEM.

**Extended Data Figure 8.**
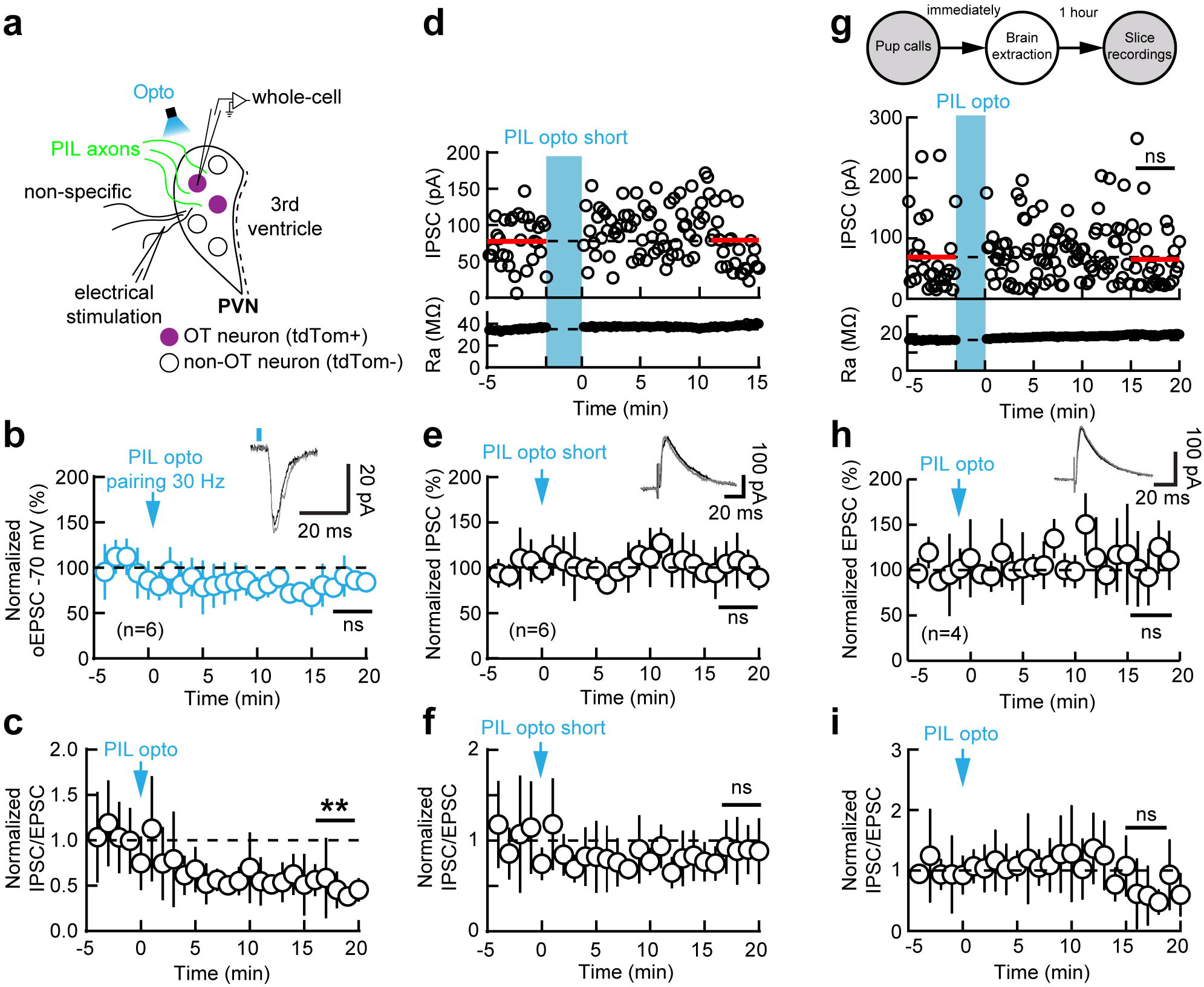
Prolonged but not brief optogenetic stimulation of PIL axons triggers iLTD. (a) Whole-cell recordings from tdTomato+ oxytocin neurons in PVN slices, optogenetic stimulation of PIL axons and placement of the extracellular stimulation electrode. (b) Whole-cell voltage-clamp recordings showing that the amplitude of oEPSCs triggered by single pulse of optogenetic stimulation was not modified following repeated optogenetic stimulation of PIL axons (‘PIL opto’; n=6 neurons, p=0.13, one-sample two-tailed Student’s t-test). (c) IPSC/EPSC ratio decreased following PIL opto (p=0.0018, one-sample Student’s t-test). (**d-f**) Repeated but brief optogenetic stimulation of PIL axons (‘PIL opto short’) did not induce iLTD: example cell (**d**; p=0.74, Mann-Whitney two-tailed test) and summary (**e**; n=6, p=0.99, one-sample Student’s t-test). (**f**) No change in IPSC/EPSC ratio following PIL opto short (p=0.49, one-sample Student’s t-test). (**g-i**) In vivo exposed to pup calls playback occlude iLTD. (**g**) Schematic of experimental protocol: example cell (**g**; p=0.9445, Mann-Whitney) and summary (**h**; n=4, p=0.9750, one-sample Student’s t-test). (**i**) No change in IPSC/EPSC ratio following PIL opto in slices of dams exposed to pup calls in vivo (p=0.7555, one-sample Student’s t-test). Data reported as mean±SD. **, p<0.01. ns: not significant.

**Extended Data Figure 9.**
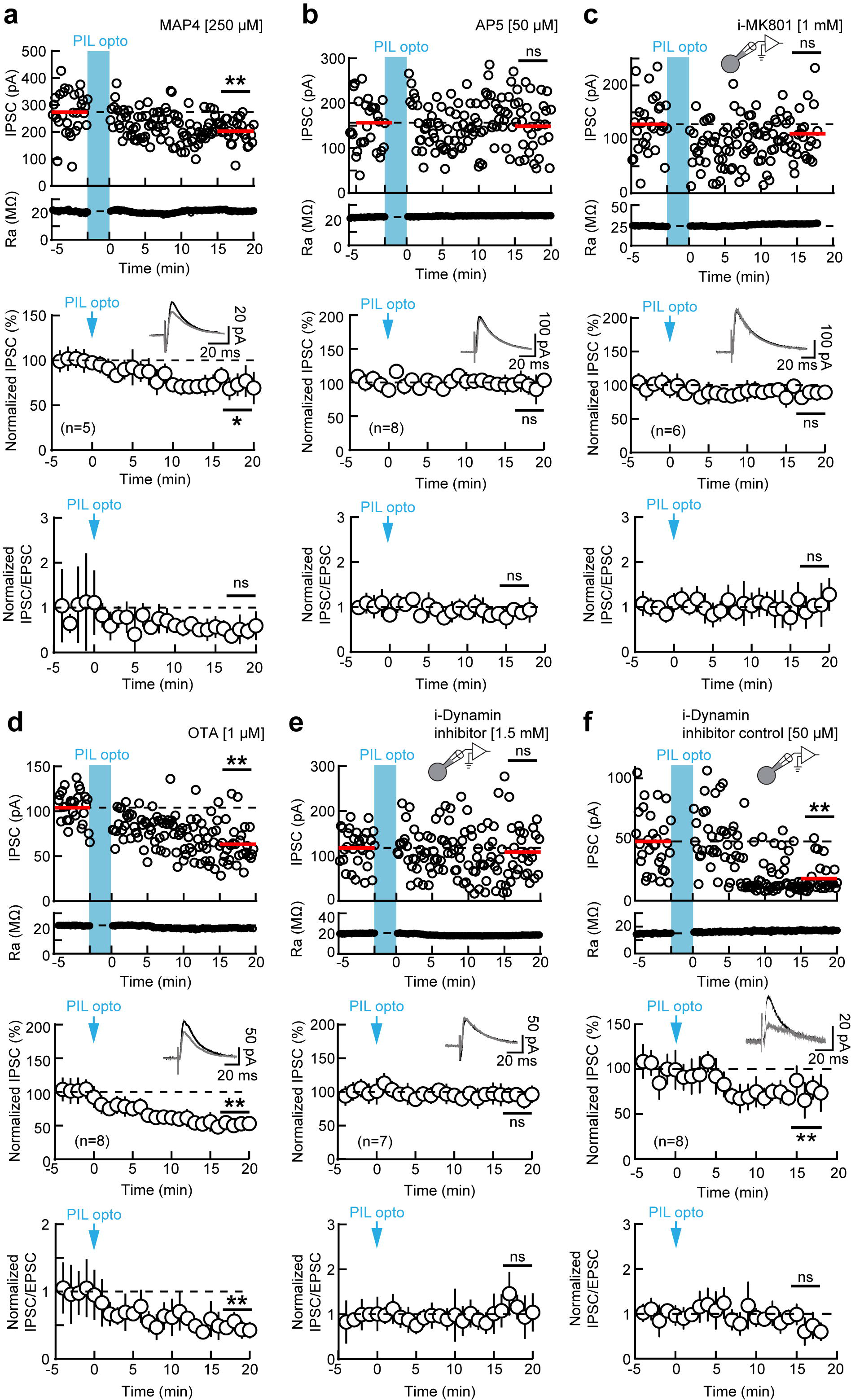
iLTD in oxytocin neurons relies on postsynaptic NMDARs and dynamin signaling. (a) Whole-cell voltage-clamp recordings showing intact iLTD after repeated optogenetic stimulation of PIL terminals in PVN (‘PIL opto’) in presence of bath-applied type-III mGluR antagonist MAP4 (250 µM), for example neuron (upper panel; p<0.0001, Mann-Whitney two-tailed test) and summary (middle panel; n=5 neurons, p=0.02, one-sample two-tailed Student’s t-test). (Lower panel) IPSC/EPSC ratio in the presence of MAP4. (b) iLTD in oxytocin neurons is NMDAR-dependent. Whole-cell voltage-clamp recordings showing no plasticity after PIL opto in presence of bath-applied AP5 (50 µM), for example neuron (upper panel; p=0.62, Mann-Whitney) and summary (middle panel; n=8, p=0.44, one-sample Student’s t-test). (Lower panel) Unchanged IPSC/EPSC ratio in the presence of AP5. (c) iLTD in oxytocin neurons is dependent on postsynaptic NMDARs. Whole-cell voltage-clamp recordings showing no plasticity after PIL opto when i-MK801 (1 mM) was applied in the recording pipette, for example neuron (upper panel; p=0.11, Mann-Whitney) and summary (middle panel; n=6, p=0.18, one-sample Student’s t-test). (Lower panel) Unchanged IPSC/EPSC ratio in the presence of i-MK801. (d) Whole-cell voltage-clamp recordings showing intact iLTD after PIL opto in presence of bath-applied OXTR antagonist OTA (1 µM), for example neuron (upper panel; p<0.0001, Mann-Whitney) and summary (middle panel; n=8, p=0.0008, one-sample Student’s t-test). (Lower panel) IPSC/EPSC ratio in the presence of OTA. (**e and f**) iLTD in oxytocin neurons is dependent on dynamin signaling. (**e**) Whole-cell voltage-clamp recordings showing no plasticity after PIL opto when i-Dynamin inhibitor (1.5 mM) was applied in the recording pipette, for example neuron (upper panel; p=0.30, Mann-Whitney) and summary (middle panel; n=7, p=0.63, one-sample Student’s t-test). (Lower panel) Unchanged IPSC/EPSC ratio in the presence of i-Dynamin inhibitor. (**f**) Whole-cell voltage-clamp recordings showing intact iLTD in oxytocin cells in the presence of a scrambled dynamin inhibitor in the recording pipette, for example neuron (upper panel; p<0.0001, Mann-Whitney) and summary (middle panel; n=8, p=0.0096, one-sample Student’s t-test). (Lower panel) IPSC/EPSC ratio in the presence of a scrambled dynamin inhibitor. Data reported as mean±SD. *p<0.05, **p<0.01, ns: not significant.

**Extended Data Figure 10.**
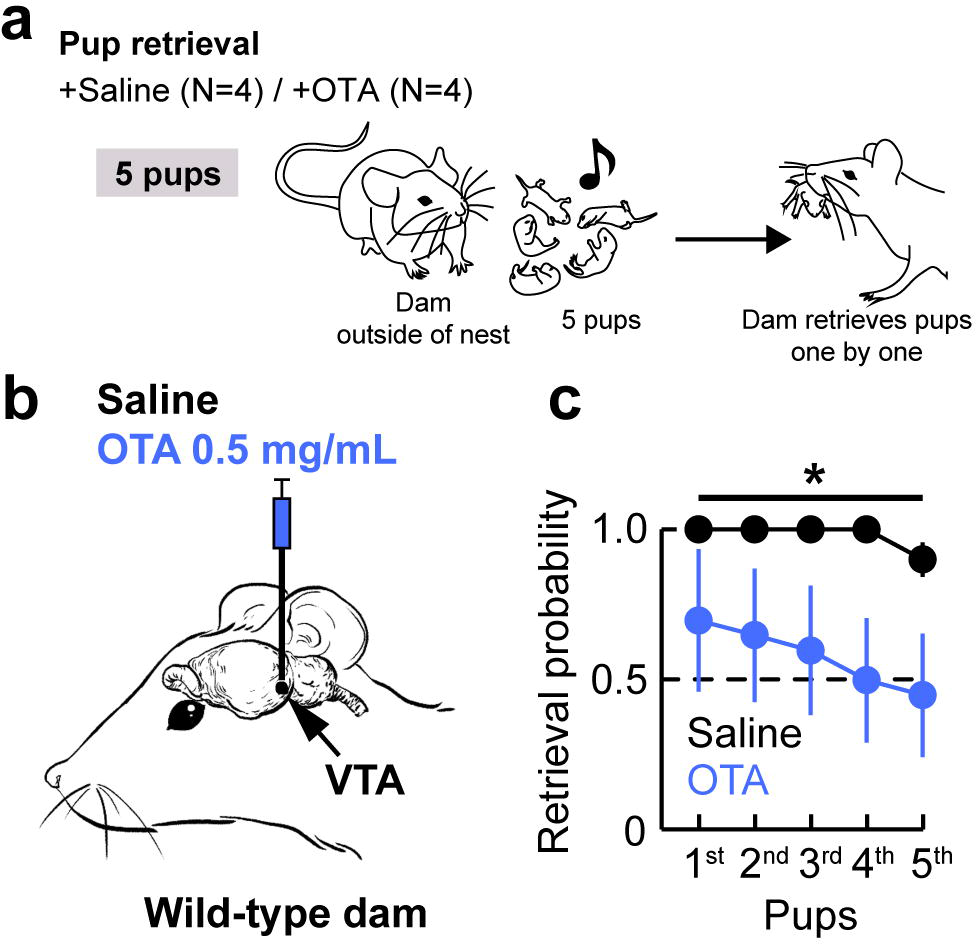
Inhibiting oxytocin signaling in the VTA impairs pup retrieval behavior. (a) Pup retrieval protocol. (**b and c**) Wild-type dams infused with the OXTR antagonist OTA (0.5 mg/mL; **b**) retrieved less pups compared to saline controls (**c**; N=4, p<0.0202, two-tailed Fisher’s test). Data reported as mean±SEM. *p<0.05.

## Notes

### Competing Interest Statement

The authors have declared no competing interest.

### Summary of Updates

New data.

